# Widespread effects of DNA methylation and intra-motif dependencies revealed by novel transcription factor binding models

**DOI:** 10.1101/2020.10.21.348193

**Authors:** Jan Grau, Florian Schmidt, Marcel H. Schulz

## Abstract

Several studies suggested that transcription factor (TF) binding to DNA may be impaired or enhanced by DNA methylation. We present MeDeMo, a toolbox for TF motif analysis that combines information about DNA methylation with models capturing intra-motif dependencies. In a large-scale study using ChIP-seq data for 335 TFs, we identify novel TFs that are affected by DNA methylation. Overall, we find that CpG methylation decreases the likelihood of binding for the majority of TFs. For a considerable subset of TFs, we show that intra-motif dependencies are pivotal for accurately modelling the impact of DNA methylation on TF binding.

## BACKGROUND

Transcription Factors (TFs) are essential regulatory proteins with diverse roles in transcriptional regulation, such as chromatin remodelling or the initiation of transcription (1). Hence, a key step to improve our understanding of the function of TFs is to identify the genomic location of TF binding sites (TFBS). It was shown that TFs usually bind to accessible chromatin (2) and therefore a variety of computational methods (3) has been developed to combine chromatin accessibility data (e.g. DNase1-seq, ATAC-seq, NOMe-seq) with TF motif information as encoded in Position Weight Matrices (PWMs) (4–7) to elucidate the tissue-specific binding profiles of TFs. Recently, LSlim-models, which capture intra-motif dependencies, have been successfully applied to overcome the nucleotide independence assumption of PWMs (8). Further approaches that allow for intra-motif dependencies include improved energy models (9), transcription factor flexible models (10), parsimonious Markov models (11), and Bayesian Markov models (12).

To provide the community with a systematic comparison of the plethora of TFBS prediction approaches, the *ENCODE-DREAM in vivo Transcription Factor binding site prediction challenge* (13) was conducted in 2016. The competing methods considered, aside from epigenomics data, also DNA shape, sequence conservation, and/or sequence composition. Interestingly, the median area under the precision recall curve (AUC-PR) for one of the winning methods across all classifiers is only 0.4 (14), suggesting that important molecular signatures influencing TF binding are not incorporated yet.

One of those signatures is DNA methylation in a CpG context. The analysis of DNA methylation has been a major focus of epigenomics research and several experimental approaches have been proposed to characterize DNA methylation *in vivo* (15): While early methods used methylation sensitive restriction enzymes in PCR and gel-based approaches (16), the usage of microarrays allowed a scale-up of CpG methylation analysis (17). Array-based methods are nowadays used to characterize the methylation levels of pre-selected CpGs, e.g. for diagnostic purposes (18). With the advancements of next-generation sequencing, several sequencing based approaches to characterize DNA methylation on a genome-wide scale have been proposed (19, 20). Most techniques used currently require bisulfite-treated DNA as input. Bisulfite treatment causes unmethylated cytosines to be converted to uracils, whereas methylated cytosines remain unchanged (21). Large-scale bisulfite sequencing studies have been performed by several international consortia such as Blueprint, Roadmap and ENCODE, to generate DNA methylation data for several tissue and primary cell types.

DNA methylation in a CpG context has been reported previously to have a repressive effect on TF binding (22). Additional studies using protein binding microarrays (23), DAP-seq (24) or methylation-sensitive systematic evolution of ligands by exponential enrichment (SELEX) (25) indicated that DNA methylation can also promote TF binding. Functionally, the addition of a methyl group to cytosines mimics a thymine and influences the steric and hydrophobic environment (26), thus called *thymine mimicry* (27). Specifically, CpG methylation leads to a widening of the major groove and narrows the minor groove (28, 29). It also affects roll and propeller twist and results in an increase of helix stiffness (29).

As summarized in (26), there are two modes how TFs can recognize DNA methylation: i) the 5 methyl-cytosinearginine-guanine triad detection and ii) the presence of van der Waals interactions between the methyl group of the cytosine and methyl groups of hydrophobic amino acids or methylene groups of polarized amino acids.

Methylation dependence has been studied in depth for several TFs such as KLF4 (30), P53 (31), CEBP complexes (25), NRF1 (32) and ZFP57 (33). However, methods specifically designed to include information about DNA methylation into the *de novo* discovery of binding motifs are rare. The mEpigram (34) software is an extension of the Epigram algorithm for motif detection (35). mEpigram derives motifs by constructing PWMs considering a sequence set derived from TF ChIP-seq data. Specifically, mEpigram computes the most enriched k-mers within the ChIP-seq peak regions compared to a randomly shuffled set of sequences. These kmers are treated as ‘seeds’ and subsequently extended both up and downstream. To incorporate DNA methylation in this process, the alphabet considered in PWM construction has been extended with a separate symbol for methylated cytosine. Viner *et al*. (36) use an alphabet with additional symbols for differently methylated cytosines and further symbols for the corresponding guanines on the opposite strand. *De novo* motif discovery is then performed by an enhanced version of the MEME suite. To analyse data generated by the *Methyl-Spec-seq* assay, Zuo *et al*. (33) use a similar extended 6-letter alphabet for PWM construction with separate symbols for methylated cytosines and guanines opposite of methylated cytosines.

Recently, the MethMotif database, which combines TF motifs with associated DNA methylation profiles, has been made available (37). In MethMotif, occurrences of known TF motifs are detected with Centrimo in ChIP-seq data from ENCODE. Subsequently, the genomic loci that are enriched for the tested motifs are overlayed with CpG methylation data from GEO. The found motifs and the CpG methylation signatures are visualized in so called *MethMotif* logos. A possible demerit of the approach pursued in MethMotif, compared with those mentioned previously, is that the methylation dependence has not been incorporated into the discovery of the TF motif.

Although the aforementioned methods demonstrated significant advantages in the characterization of TF binding sites by including DNA methylation, they do suffer from the simplifying *independence of nucleotide assumption* made in PWM models. Even without considering DNA methylation, several recent studies demonstrated that including intra-motif dependencies improves the accuracy of motif models. The models employed for this purpose include variable-order Bayesian networks (38), Bayesian Markov models (12), transcription factor flexible models (10), parsimonious Markov models (11, 39), and sparse local inhomogeneous mixture (Slim) models (8). Considering DNA methylation, the independence assumption is obviously violated in a CpG methylation context. In addition, neither MethMotif nor mEpigram provide the user with means to perform methylation-aware genome wide TFBS predictions.

Here, we present MeDeMo (Methylation and Dependencies in Motifs), a toolbox using an extension of Slim models capturing intra-motif dependencies, which accounts for the presence of DNA methylation. We illustrate that the combination of methylation information and intra-motif dependencies considered by MeDeMo typically yields an improved prediction performance compared with a standard PWM-based approach. To this end, we analysed the DNA methylation dependence of hundreds of TFs in cell-lines and primary cells using DEEP and ENCODE data. MeDeMo is available as a stand-alone tool allowing both the inference of methylation-aware TF motifs and to obtain genome-wide TFBS predictions.

## RESULTS AND DISCUSSION

To test whether the inclusion of cell type-specific methylation information and explicitly modelling dependencies within DNA-binding sites is beneficial for a specific TF, we follow the procedure illustrated in Fig. 1. We start from whole-genome bisulfite sequencing data for the cell type at hand, discretize methylation calls by the betamix (40) approach, and use these binary methylation calls to convert the original hg38 genome sequence into a methylation-aware genome version. Specifically, we convert methylated ‘C’ to ‘M’ and ‘G’ opposite of a methylated ‘C’ to ‘H’, yielding an extended 6-letter alphabet.

**Fig. 1.**
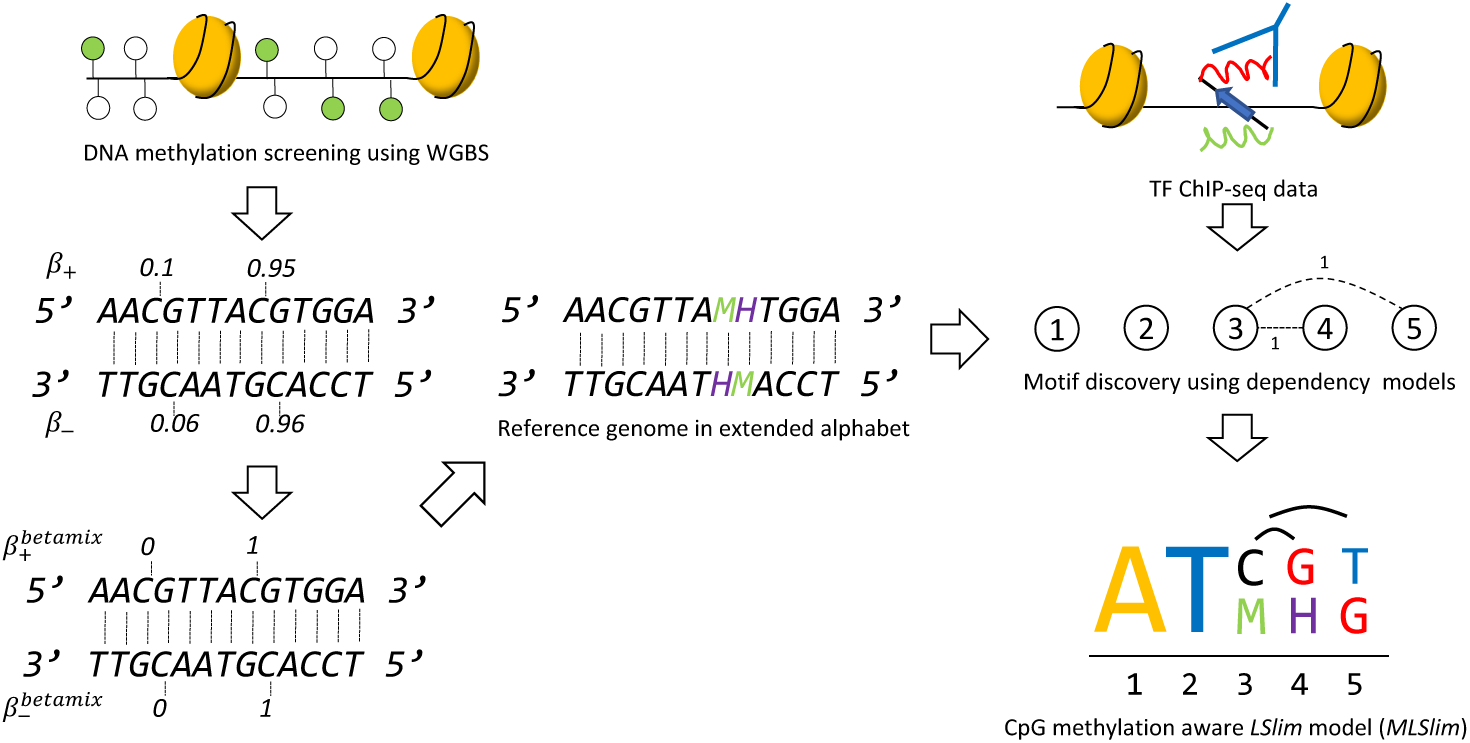
Overview of the MeDeMo workflow: (1) DNA methylation is assessed using whole genome bisulfite sequencing. (2) DNA methylation is quantified using *β*-values. (3) Methylation calls (*β*-values) are discretised using the betamix approach resulting in a binary methylation state for each Cytosine in a CpG context. (4) A novel reference genome is generated by denoting occurrences of methylated cytosines with the letter M and occurrences of guanines opposite of a methylated cytosine with the letter H. (5) In-vivo transcription factor binding site information are obtained using peak calls from TF -ChIP-seq data. (6) TF binding data is used for motif discovery with LSlim models on the methylation aware reference genomes; (7) resulting in methylation aware TF motif representations.

Based on the ChIP-seq peaks downloaded from ENCODE, we extract sequences under the peaks, which serve as input to the de novo motif discovery. As statistical binding site models, we use either Position Weight Matrix (PWM) (41, 42) assuming independence of nucleotides, or LSim(5) (8) models capturing dependencies between nucleotides over a distance of at most 5 nucleotides. Both types of models are applied to sequences under peaks extracted from the original hg38 genome, or to sequences under peaks extracted from the methylation-aware genome version for the cell type of the ChIP-seq experiment. This results in four modelling alternatives (Table 1), namely i) PWM applied to original hg38, ii) PWM applied to the methylation-aware genome, iii) LSlim applied to original hg38, and iv) LSlim applied to the methylation-aware genome.

**Table 1.**
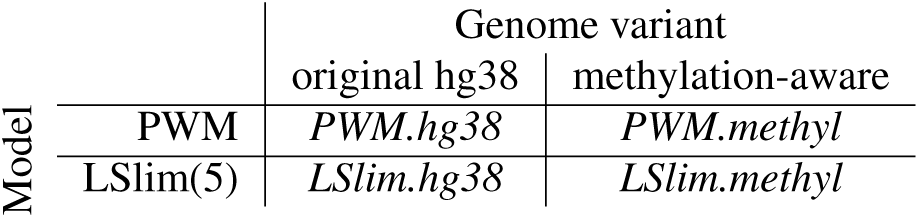
Overview of the combinations of genome variants and motif models considered in this study.

In the remainder of this section, we first investigate for which TFs the introduction of a methylation-aware genome and the inclusion of dependencies yield an improvement in classification performance discriminating bound from unbound sequences. We then consider specific examples of TFs that show such an improvement, discuss their binding motifs in relationship to methylation, and study general trends in sensitivity of TFs to methylation of their binding sites. We finally present prototypical examples of TFs for which the combination of methylation information and modelling dependencies is pivotal to optimal performance.

### Investigating the impact of DNA methylation on binding

For benchmarking the different modelling alternatives, we follow a classification-based approach. Here, motif models are tested for their capability of distinguishing bound from unbound sequences. We consider as sequences bound by a specific TF those under a ChIP-seq peak, whereas unbound sequences sampled uniformly across the genome (cf. Methods). Since for the majority of TFs, this is a highly imbalanced classification problem, we use the area under the precision-recall curve (43) as a performance measure. For each TF, we collect all data sets that are available from ENCODE for the cell types under study (GM12878, HepG2, K562, liver), which might include replicate experiments for the same combination of cell type and TF, e.g., performed in different labs.

We further follow a 10-fold cross validation strategy to be able to also assess classification performance on the data from the same experiment. For each partition of the 10-fold cross validation, we consider the motif reported on rank 1 by the SlimDimont framework during training (cf. section Training procedure) for evaluating model performance on test data.

In the following, we distinguish *within* cell type (i.e., training and test cell types match) and *across* cell type (i.e., training and test cell types are different) classification performance. For each of these sub-sets of classification problems, we collect all AUC-PR values and perform a one-sided Prentice rank sum test (44, 45) (using `prentice.test` from R-package `muStat`) between each pair of modelling alternatives considering cross-validation folds as replicates of the same experiment (replicated block design) and using a significance level of *α* = 0.05. In addition, we count the number of data sets, for which one alternative yielded a higher classification performance than the second one. Finally, we visualize the differences of AUC-PR values in violin plots as shown in Fig. 2.

**Fig. 2.**
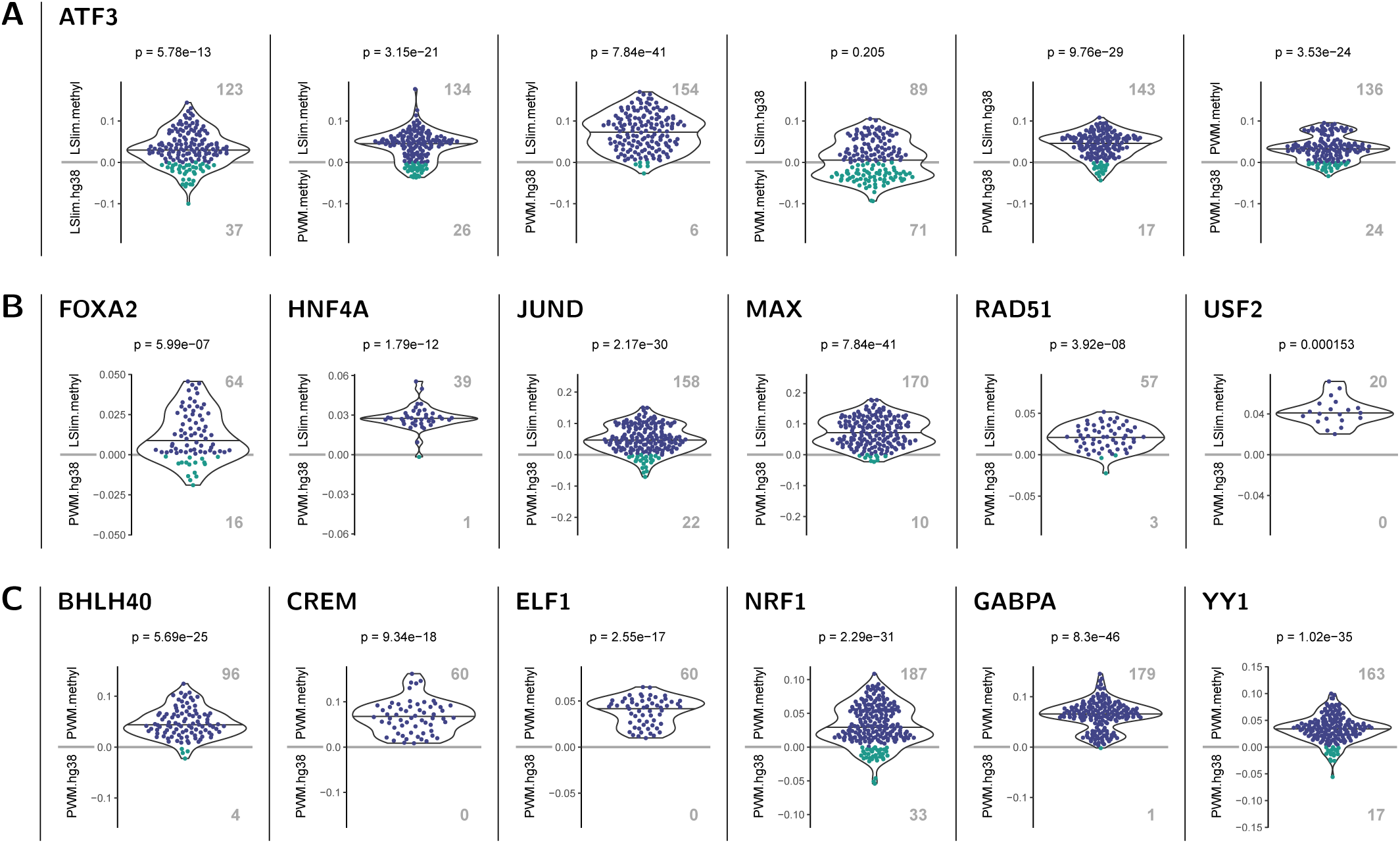
Examples of TFs with significantly improved classification performance (AUC-PR) in across cell type predictions using a methylation-aware genome. Each panel shows a pairwise comparison of models as indicated by the y-labels above and below the zero line. Each dot represents a case (data sets × cross validation folds) with different colours for positive (i.e., top model performs best) and negative (i.e., bottom model performs best) differences of AUC-PR values. Total number of cases where one model performs better than the other are shown as boldface, grey numbers. In addition, points are summarised by a violin plot and corrected p-values for the *H*_0_ that both models perform identical (Prentice test) are given in the header. (A) Pairwise comparison of different modelling variants for ATF3. We find that all methylation-aware models perform better than their counterparts learned on the original hg38 genome and that dependency models (LSlim) perform better than PWM models on the same genome variant. For instance, LSlim.methyl performs better than LSlim.hg38 in 123 cases, whereas the opposite is true for only 37 cases, leading to a p-value of 5.78 *×* 10^−13^. (B) Comparison of methylation-aware dependency models (LSlim.methyl) with PWM models using standard hg38 (PWM.hg38) for TFs with a clear advantage of the combination of methylation information and modelling dependencies. (C) Comparison of PWM models learned from the methylation-aware genome with those learned from the standard hg38 genome.

Analysis of binding models for ATF3 (across cell type setting) is presented as an example in Fig. 2A. We show the corresponding results for all six pairwise comparisons of the four modelling alternatives. For instance, from the leftmost panel of Fig. 2A, we observe that the difference between the AUC-PR values of LSlim.methyl and LSlim.hg38 are mostly positive indicating an improved performance of LSlim models on the methylation-aware genome compared with standard hg38. This difference is statistically significant with a p-value of 5.78 × 10^−13^, where for 123 cases (data sets × cross validation folds) LSlim.methyl performs better than LSlim.hg38, whereas the opposite holds for only 37 cases. Similarly, we find a significant improvement of LSlim.methyl over PWM.methyl (indicating that dependencies are beneficial), of LSlim.methyl over PWM.hg38, of LSlim.hg38 over PWM.hg38 and of PWM.methyl over PWM.hg38. For the comparison of LSlim.hg38 (only dependencies) with PWM.methyl (only methylation information), we do not observe a significant difference, which indicates that both aspects of the novel approach contribute to a similar degree to the final classification performance of LSlim.methyl. Together, these results make ATF3 a prototypical example of a TF for which the combination of methylation information and modelling dependencies is important for yielding the best classification performance among the considered classification approaches.

In Fig. 2B, we present further examples of TFs for which the combination of methylation information and modelling dependencies is beneficial. These cases also illustrate the varying quantity of combinations of training and test data sets from different cell types available for different TFs (each split into 10 cross validation folds). Here, these span from 2 (USF2, one ChIP-seq data set for each of two cell types) to 18 (JUND and MAX). In all cases, the improvement of LSlim.methyl over PWM.hg38 is statistically significant, although the magnitude of the improvement in classification performance (y-axis) as well as the proportion of cases where one model performs better than the other differ among these TFs.

Fig. 2C shows examples of TFs for which the improvement of PWM.methyl over PWM.hg38 is significant but the improvement of LSlim.methyl over PWM.methyl is not, i.e., TFs for which inclusion of methylation information is beneficial but modelling dependencies does not lead to further improvements.

### TFs showing sensitivity to DNA methylation

We compile an overview of such pairwise comparisons of modelling alternatives in Fig. 3. Here, we apply stringent criteria for counting one modelling alternative to perform better than a second one for the TF at hand. Specifically, we require the improvement to be significant i) in the within cell type *and* across cell type settings consistently for both training variants (shuffled and randomly drawn negatives, cf. Methods).

**Fig. 3.**
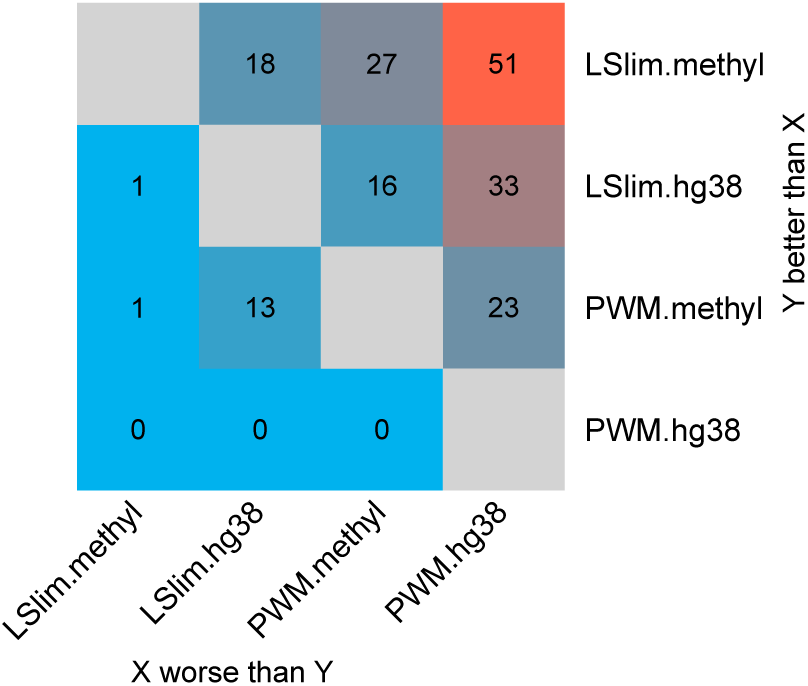
Pairwise comparison of different modelling variants. For each pair of models (PWM, LSlim) and each genome variant (original: hg38, methylation aware: methyl), we determine the number of TFs for which the model listed in the row performs significantly better than the model listed in the column i) within and across cell types, and ii) consistently using randomly drawn and shuffled negatives.

Of the 335 TFs considered in total, ChIP-seq data sets for at least two cell types are available for 143 TFs, while all remaining cannot meet the stringent criteria by definition.

Among these 143 TFs, we observe the largest number of TFs (51) with significant and consistent improvement comparing LSlim.methyl (methylation information *and* dependencies) against PWM.hg38 (neither of the two). We also find improvements for a substantial number of TFs when considering intra-motif dependencies in addition to methylation information (i.e., LSlim.methyl compared with PWM.methyl, 27 TFs), or when considering methylation information in addition to intra-motif dependencies (i.e., LSlim.methyl compared with LSlim.hg38, 18 TFs). Modeling only dependencies (LSlim.hg38 vs. PWM.hg38) or including only methylation information (PWM.methyl vs. PWM.hg38) yields an improvement for 33 and 23 TFs, respectively. For the direct comparison of either including only dependencies (LSlim.hg38) or only using a methylation-aware genome (PWM.methyl), we find balanced numbers of TFs with an improvement in either direction (16 and 13 TFs). The opposite comparisons yield a significant improvement only for a minority of at most one TF. Considering the traditionally used PWM model using the standard hg38 genome, we find a better performance for PWM.hg38 compared with LSlim.hg38, LSlim.methyl or PWM.methyl for none of the TFs studied. We find one TF (HDAC2) for which the PWM model yields a better performance than the LSlim model on the methylation-aware genome. In this case, the PWM.methyl model significantly outperforms all other modelling alternatives and adding dependencies appear to be rather detrimental. We further find one TF (CTCF) for which the LSlim model works better on the original than on the methylation-aware genome. Converse to HDAC2, intra-motif dependencies seem to be of greater importance for CTCF than methylation information, and the LSlim.hg38 outperforms any other modelling alternative.

The examples previously shown in Fig. 2A/B are in the intersection of all three sets for which LSlim.methyl performs better than any of the other three alternatives (top row of Fig. 3), whereas those shown in Fig. 2C are from the union of LSlim.methyl vs. LSlim.hg38, LSlim.methyl vs. PWM.hg38 and PWM.methyl vs. PWM.hg38, excluding TFs where one model on the original hg38 genome performs better than its methylation-aware counterpart.

We present a list of those TFS for which methylation information was beneficial for prediction performance in Table 2. Here, we exclude TFs without direct and sequence-specific DNA binding (as discussed for BRCA1 below), while we provide a complete list of TFs in Supplementary Table 2.

**Table 2.**
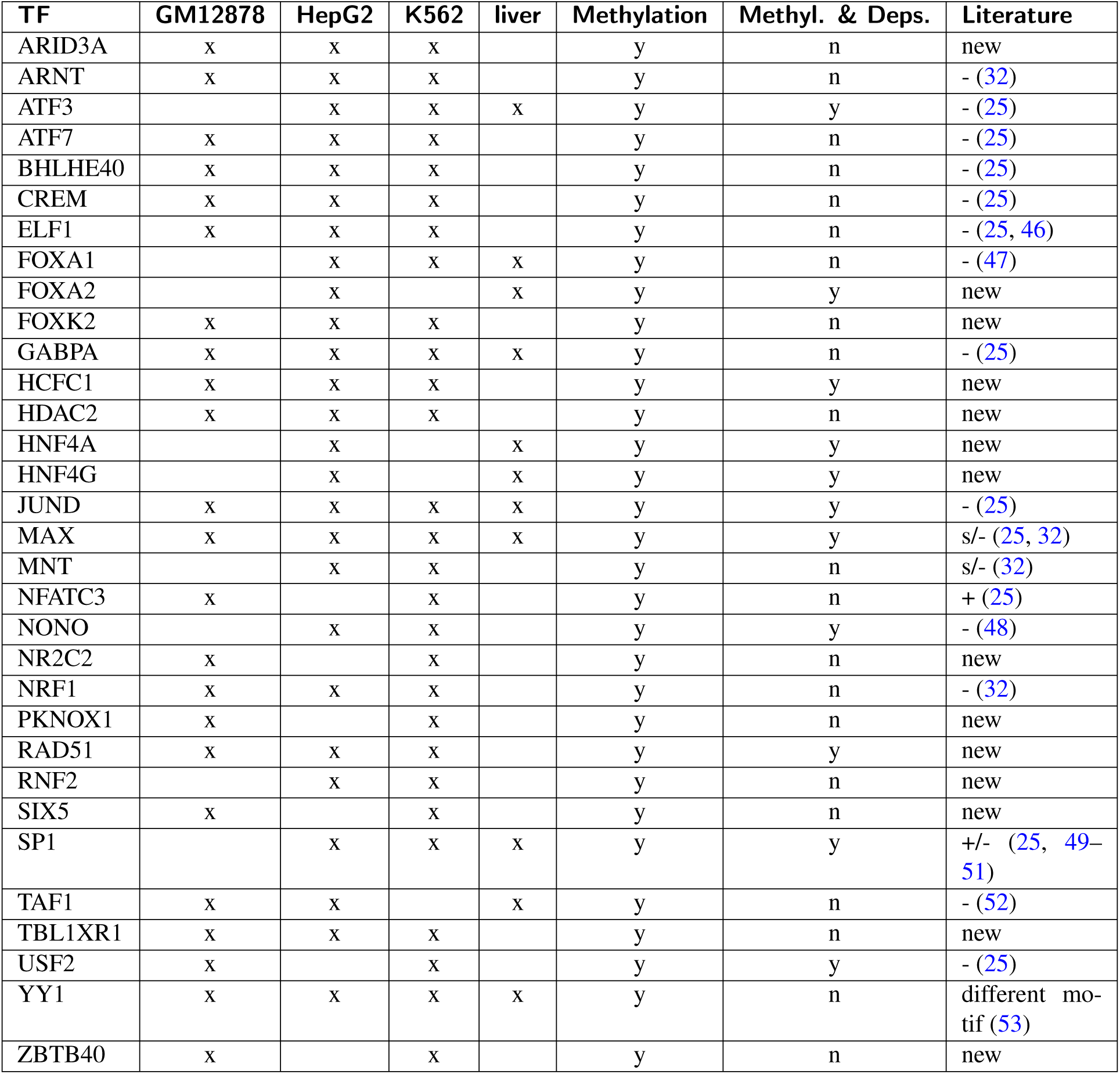
Summary of TFs that profit from considering DNA methylation in the motif models. For each TF, we list the availability of ChIP-seq data sets for the four cell types studied. Columns “Methylation” and “Methyl. & Deps.” indicate a significant and consistent improvement (y: yes, n: no) by including information about methylation sensitivity in general and/or in combination with modelling intra-motif dependencies, respectively. In the last column, we note references to the literature for TFs that have already been reported to be methylation sensitive, where “-”, “+” and “s” indicate negative or positive influence of methylation or general methylation sensitivity, respectively.

### Methylation sensitivity of TFs

Having established a set of TFs for which the inclusion of methylation information leads to an improvement in the benchmark study, we further investigate binding preferences of TFs in the context of their binding motifs. To this end, we compute a position-specific profile of methylation sensitivity by altering CpG dinucleotides within putative binding sites to their fully methylated variant MpH and recording the resulting differences in the corresponding binding scores according to the motif model. By this means, we may decode the information about methylation preference captured by the motif model. If the difference of binding scores is positive, this corresponds to MpH dinucleotides (i.e., methylated DNA) being preferred over CpG dinucleotides by the model at a given position, and vice versa. By referring to the level of predicted binding sites, this measure of methylation sensitivity is easily transferred to LSlim models, where methylation sensitivity may depend on the sequence context.

In Fig. 4, we present six examples of such profiles of methylation sensitivity according to the corresponding PWM models, plotted below the sequence logo of their predicted binding sites. As might be expected, all these examples have in common that their motifs contain prominent CpG dinucleotides, although with different frequencies and in different contexts. For ELF1, CREM, and MAX, we observe one prominent CpG dinucleotide as part of their motifs, where CpG content varies between 0.57 (ELF1) and 0.85 (CREM). In all three cases, methylation of this CpG dinucleotide according to the model leads to a decrease in the prediction score, indicating that methylation is detrimental for binding affinity. Similar patterns also occur for YY1 with one prominent and several less frequent CpG positions, and for BRCA1 and NRF1 exhibiting two prominent CpG dinucleotides each.

**Fig. 4.**
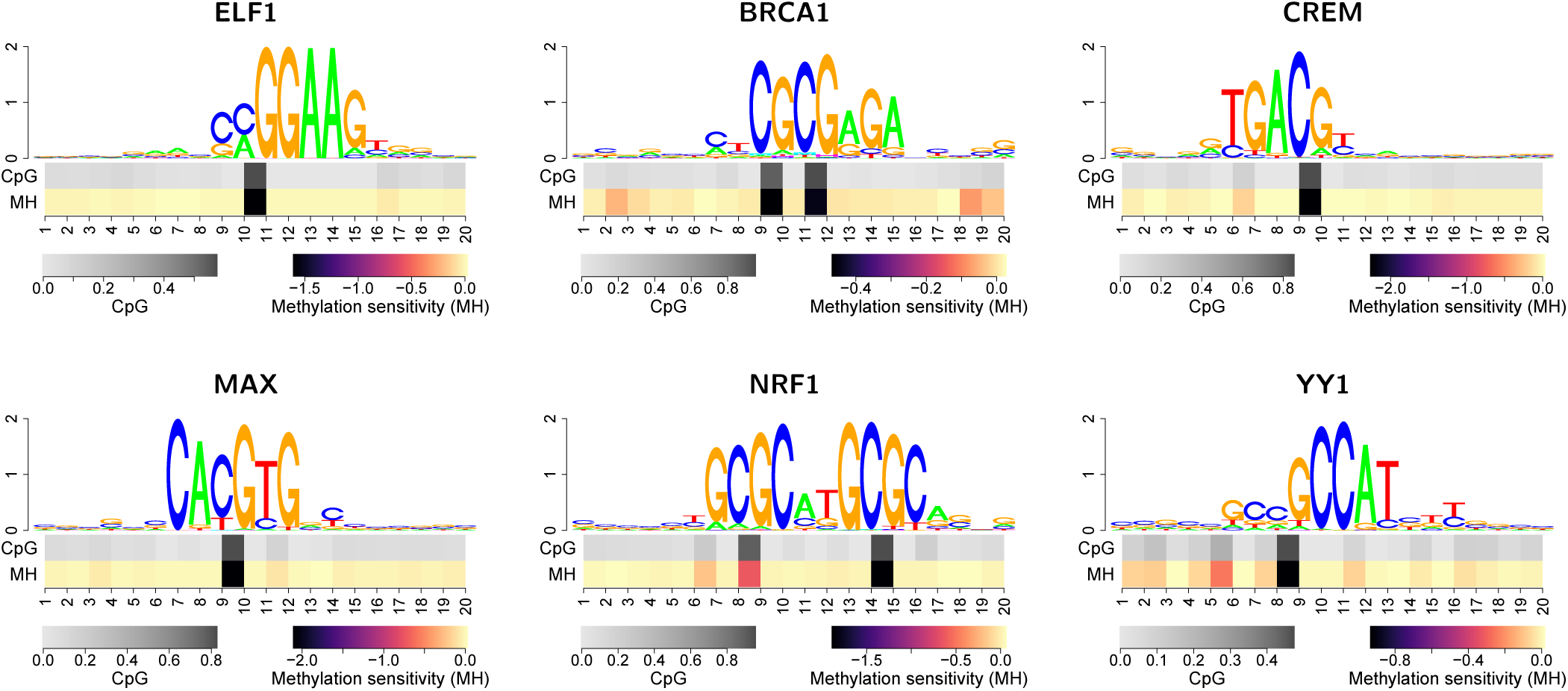
Methylation sensitivity of TFs with improved performance using a methylation-aware genome. In each panel, the top part show a sequence logo of the discovered motif using the extended alphabet. However, since the model learned to penalize methylated DNA in all six cases, additional symbols are only visible in case of BRCA1. In the bottom part of the plot, we visualize position-specific CpG content (top row with grey scale) and methylation sensitivity (bottom row with colour scale) within predicted binding sites. Positive values of methylation sensitivity indicate preferred binding of methylated DNA, whereas negative values indicate methylated DNA being disfavored. For all six TFs, we observe a detrimental effect of DNA methylation at frequent CpG positions.

For NRF1, it appears as if methylation affects one of the CpGs (position 8/9) to a lesser degree than the other (position 14/15). However, ChIP-seq does not provide strand information and the strand model encapsulating the PWM allows for switching the strand orientation of the binding site. For these reasons, and because the motif of NRF1 is clearly palindromic, this phenomenon needs to be interpreted with care. An alternative explanation might be that once one of the CpGs present in NRF1 binding sites is methylated, additional methylation of the other CpG does not lead to a substantial further effect. Notably, the binding motif discovered for BRCA1 does not match the canonical motif present in HOCOMOCO (54). BRCA1 has been reported to bind DNA directly but without sequence specificity (55). The ZBTB33-like motif discovered by our approach could possibly be due to indirect binding, and a similar motif has been reported for BRCA1 before (56).

Strikingly, the influence of methylation on the prediction score at high-CpG positions is negative in all examples presented in Fig. 4, suggesting that DNA methylation may lead to reduced binding affinity for many TFs. In order to investigate if this observation constitutes a general tendency among the studied TFs, we consider all TFs with a significant and consistent improvement in prediction performance when including methylation information (cf. Table 2). For each of these TFs, we compute the corresponding profiles of methylation sensitivity per data sets and record the range of values (i.e., minimum value to maximum value) present in the profile. Strong deviations from 0 of the maximum or minimum value indicate a clear preference for methylated or unmethylated DNA according to the model, respectively. We find (Fig. 5) that the maximum value is only slightly above 0 for the wide majority of TFs, whereas for many TFs, the minimum value is clearly below 0. This indicates that for most TFs, the profiles of methylation sensitivity indeed are similar to those presented in Fig. 4. There are a few examples of TFs (FOXA1, FOXA2, HNF4A, HNF4G, RAD21), for which neither the maximum nor the minimum of methylation sensitivity shows a strong amplitude. These TFs do not have a prominent CpG in their binding motifs. Nonetheless, inclusion of methylation information leads to an improvement in prediction performance. We discuss possible explanations of this observation for two examples below (FoxA1 and FoxA2, Fig. 6).

**Fig. 5.**
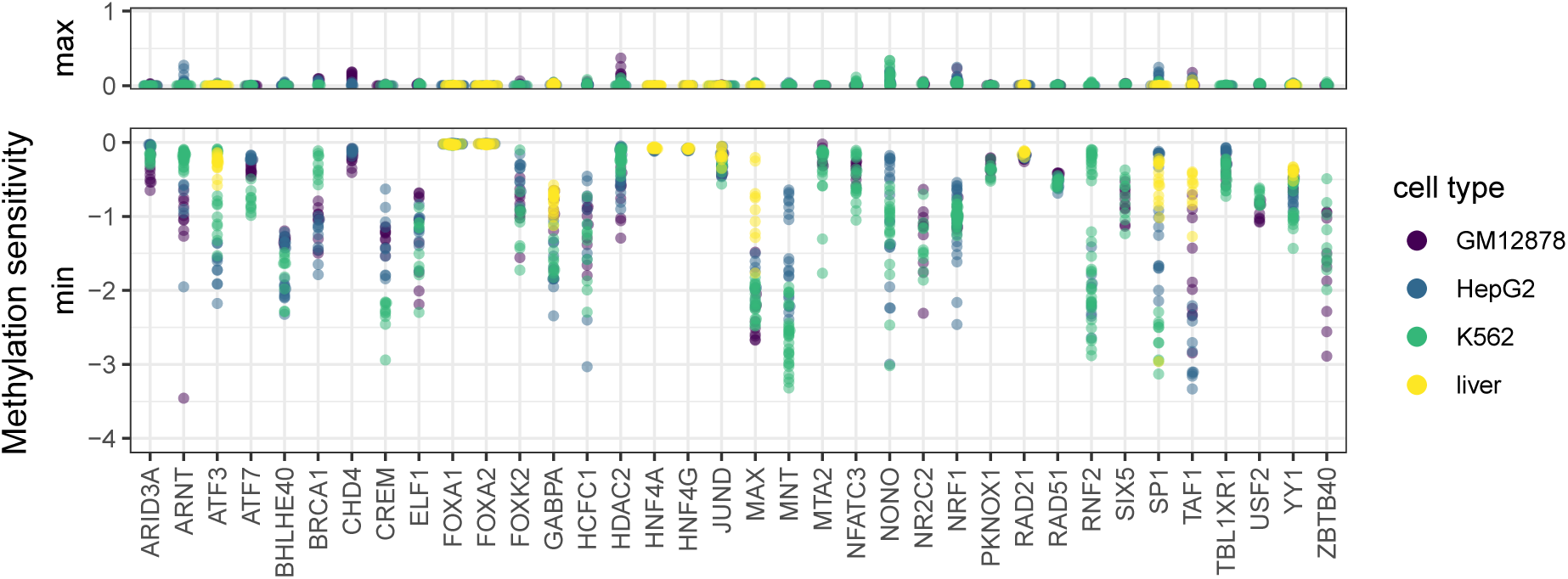
Methylation within binding motifs is mostly detrimental in models with significantly and consistently improved prediction performance (cf. Figs. 2 and 3). For each TF and each data set, we record the profiles of methylation sensitivity as shown in Fig. 4. We aggregate this profile to two values per data set by computing the minimum and maximum value of methylation sensitivity, which captures the range of values observed in the profile. Here, we plot these maximum and minimum values of methylation sensitivity across all training data sets. We observe a large amplitude of negative values for the minimum (i.e., methylated DNA being disfavored) but only slightly positive values for the maximum, indicating that – according to the models – DNA methylation is detrimental for the majority of TFs.

**Fig. 6.**
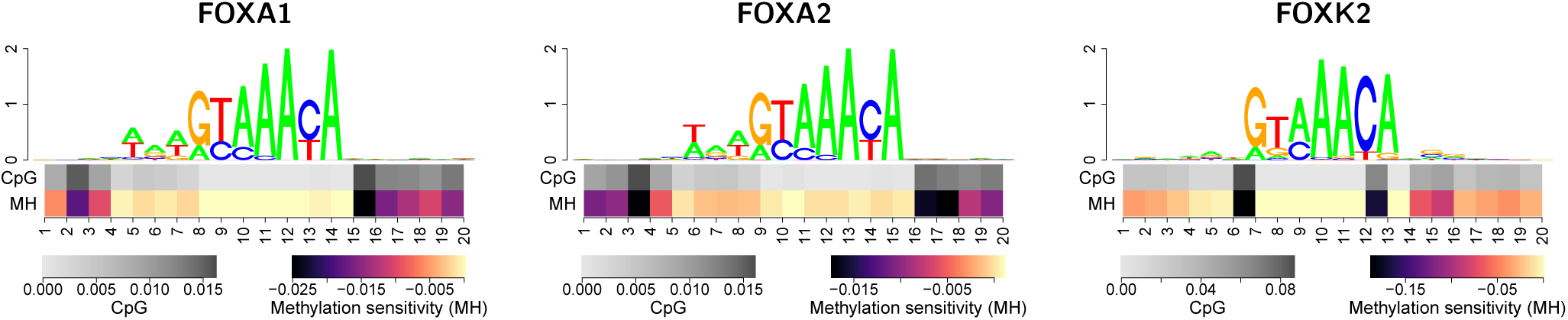
Methylation sensitivity may differ between members of a TF family. While methylation sensitivity of FOXA1 and FOXA2 is highly similar in HepG2 cells, that of FOXK2 is noticeably different, although all three motifs appear to be highly similar. This behaviour is consistent between different cell types (Suppplementary Fig. 2).

For several of the TFs in Fig. 5, a negative influence of methylation on their binding has been reported before. This includes ARNT (32), ATF3/7 (25), CREM (25), ELF1 (25, 46), GABPA (25), JUND (25), MAX (25, 32), MNT (32), NRF1 (32), USF2 (25), and YY1 (53). SP1 shows a generally negative influence of methylation on its binding sites in our data, although with cell type-specific strength. Previous results for SP1 have been contradictory, as some studies suggested a positive influence of binding site methylation (25), whereas others indicated no decisive influence (50), negative effects (49), or the prevention of methylation by SP1 binding (51). In general, preference for de-methylated DNA may be observed either due to the direct binding preference of the TF at hand, or due to a de-methylation of the bound region as an effect of TF binding. Based on our data, these two cases could not be distinguished.

The reasons for the mostly detrimental influence of methylation for the TFs in our study could be manifold. First, this could be a bias introduced by the specific selection of TFs under study, although no such bias has been introduced intentionally, since we consider all TFs with ENCODE data sets in at least two of the selected cell types. Specifically, CEBPB (27, 57, 58), SMAD5 (25) and ZBTB33 (25, 59, 60) have been reported to prefer methylated DNA, but we did not observe a significant and consistent improvement of prediction performance in our study. For GATA1/2/4 (46, 58), IRF2 (25), KLF16 (25), NFATC1 (25), STAT1/5A (46) and ZNF274 (25), we had only data for one of the cell types studied, which prevented us from studying performance across cell types. Second, this result might be an artifact of our method. While we cannot rule out this possibility in general, we do observe clearly positive methylation sensitivity values for a few TFs. Examples (ZBTB33 with inconsistent results across cell types, and NFATC1 and ZNF274 with ChIP-seq data available only for one cell type) are given in Supplementary Fig. 1. Hence, we may at least conclude that our method is capable of capturing such patterns in general. Third, there might also be a bias of methylation on the ChIP-seq experiment that constitute the basis of our approach, although we did not find this to be reported before. For instance, methylation might influence the amplification step in the ChIP-seq protocol, which could lead to an under-representation of reads from methylated peak regions.

### Methylation sensitivity may vary within a TF family

As we had ChIP-seq data from TFs with the same binding domain (family) and similar consensus sites we wondered, whether there could be differences in the sensitivity to DNA methylation for individual family members. For example, FOXA1 and FOXA2 showed a low amplitude in methylation sensitivity in Fig. 5, whereas FOXK2 binding appears to be more strongly affected by DNA methylation. Although all three TFs are members of the forkhead box family, they play different roles related to development and disease (61, 62). In Fig. 6, we present the binding motifs and profiles of methylation sensitivity discovered by our approach for FOXA1, FOXA2, and FOXK2 in HepG2 cells. In general, all three motifs follow the consensus GTAAAYA with slight deviations. The major difference between FOXA1/FOXA2 and FOXK2 motifs is an additional A/T-rich stretch directly preceding this canonical motif. With regard to methylation sensitivity, we find more prominent difference between the three TFs. Specifically, FOXA1 and FOXA2 exhibit a mildly negative effect of methylation at positions bordering their core motif. While the influence on the binding score of any of these positions individually is rather low, the combination of multiple methylated CpGs at bordering positions might still have an effect on binding site prediction. By contrast, FOXK2 shows two, still rather infrequent, CpG dinucleotides at positions 6/7 and 12/13 of the core motif, which are not present in the FOXA1/FOXA2 motifs. Both of these positions show a stronger sensitivity to methylation than any position of FOXA1/FOXA2. This general picture is consistently observed in other cell types (Supplementary Fig. 2). Biologically, this observation might be linked to the mechanism of FOXA1 and FOXA2 acting as pioneering factors (61, 63), although pioneering activity has been shown for FOXK2 as well (62).

### DNA methylation sensitivity depends on sequence context

In this study, we identified a substantial number of TFs, for which the combination of methylation information and modelling intra-motif dependencies yields an improvement in classification performance compared with the base model (PWM on original hg38) but also relative to the individual contributions of methylation information and/or modelling dependencies (cf. Figs. 2B and 3). Here, we discuss three of those TFs in more detail that illustrate the breadth of the binding landscapes observed and how these are linked to specific profiles of methylation sensitivity. In Fig. 7, we present dependency logos (8, 64) of the predicted binding sites of JUND (K562 cells), USF2 (K562), and ATF3 (HepG2), which are enriched with partition-specific profiles of methylation sensitivity.

**Fig. 7.**
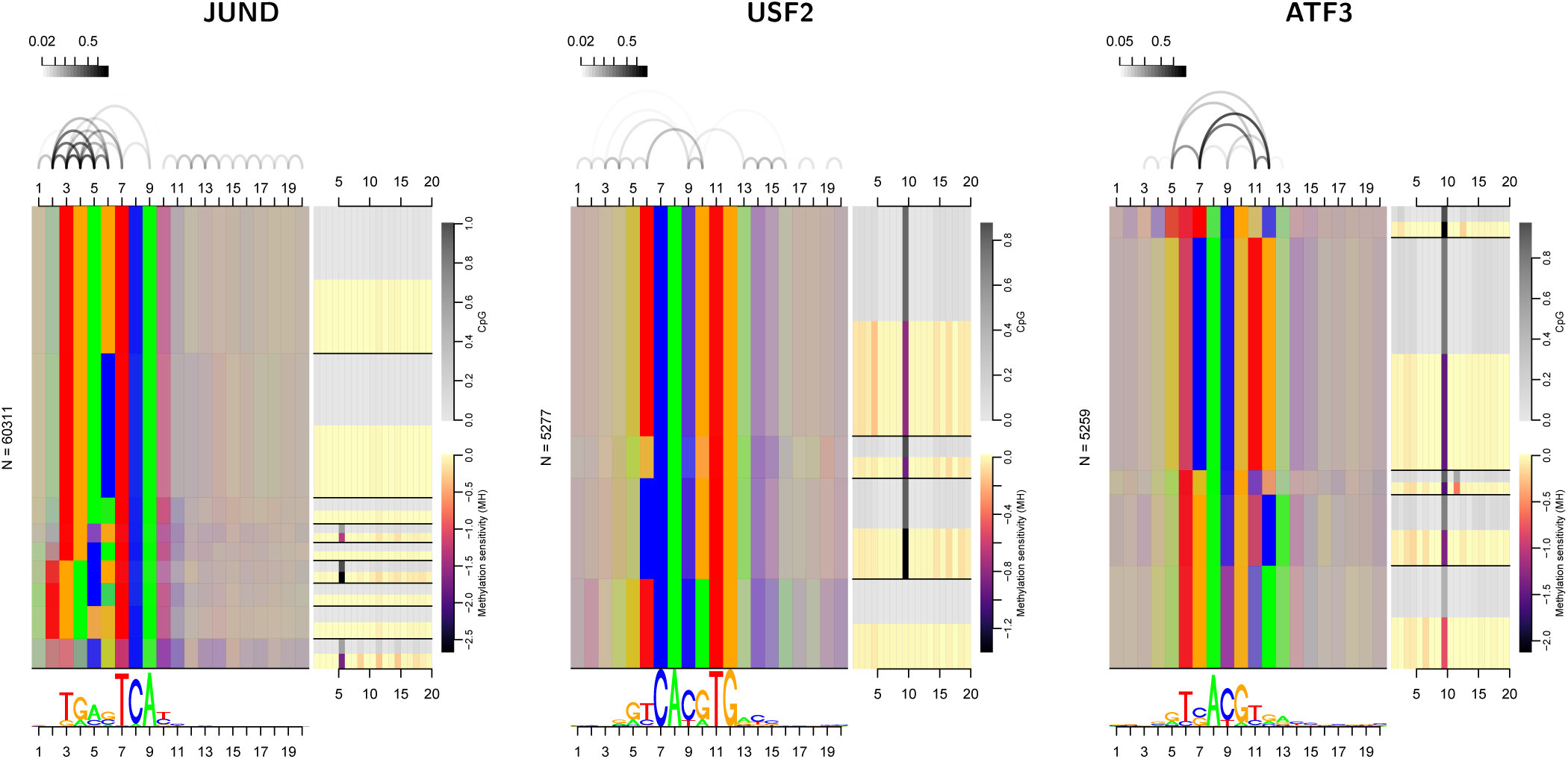
For JUND, USF2, and ATF3, the advantage of combining dependency models with a methylation-aware genome can be attributed to specific properties of the corresponding binding landscapes. For JUND, we find the known variable spacer between the two 3 bp half motif (TGA, TCA), where only the longer spacer frequently contains a CpG. For USF2, the prevalent CpG at positions 9 and 10 shows dependencies to other binding site positions, and is not present in one specific subset (TCACATG) of binding sites. For ATF3, we find broad heterogeneity, where each sub-motif contains CpG at positions 9 and 10 in different proportions.

JUND binds DNA as a dimer with a variable 1-2 bp spacer (65) which may be captured by dependency models like the LSlim model employed in this study (8) and more specialized models like TFFMs (10), but not (adequately) by standard PWM models. In the dependency logo, this variable spacer is visible as two distinct blocks, the upper block starting with consensus TGA at positions 3-5 and the lower, smaller block starting with the same consensus (TGA) but already at positions 2-4. Both variants share the consensus TCA at positions 7-9. For the short-spacer variant (upper block), only a small subset of binding sites deviating from the standard consensus (TGYGTCA, 4th partition from top) has a substantial fraction of CpG dinucleotides at positions 5/6, which are moderately methylation sensitive. By contrast, about a quarter of the long-spacer variant (lower block, 6th partition from top) with consensus TGACGTCA exhibits a CpG dinucleotide at positions 5/6, which are strongly affected by methylation. Both, the variable spacer and the specific profiles of methylation sensitivity within both variants, explain why the combination of methylation information and modelling intra-motif dependencies yields a particular advantage for JUND binding sites. Notably, the JUND motif for K562 present in the MethMotif database (37) only represents the short-spacer variant and no specific methylation profile within the core motif, where both likely is an effect of its limitation to PWM models. By contrast, our results suggest that both spacer variant and the associated patterns of methylation sensitivity are present across cell types (Supplementary Fig. 3).

For USF2, we observe a canonical E-box motif with consensus CACGTG for the majority of binding sites, and consensus CACATG for a minority of binding sites displayed as the bottom partition of the dependency logo. Intra-motif dependencies are especially prominent between positions 6 and 10, but also several positions flanking the core motif. The dependency between positions 6 and 10 can be attributed to the consensus CACATG always being preceded by a T at position 6, whereas the canonical E-box motif may also be preceded by C or G. Only those binding sites following the consensus CAYGTG frequently (approx. 80%) exhibit a CpG at positions 9/10, which is then moderately (1st and 2nd partition from top) or strongly (3rd partition from top) affected by methylation. For the partition with consensus CACATG, we find an almost flat profile of methylation sensitivity. Again, this dependency structure and associated varying methylation sensitivity may adequately be captured by dependency models but not by standard PWM models.

Finally, we observe substantial heterogeneity among the binding sites of ATF3, which have been reported before (8). Starting from the top of the dependency logo, we find a partition with consensus TTTACGRC (positions 5-12), followed by a large partition with consensus YCACRTG (positions 6-12), a small partition with consensus TRACGYR (positions 6-12), a partition with consensus TGACGBCA (positions 6-13), and finally a partition with consensus TGAYGYAA (positions 6-13). The diversity of the predicted ATF3 binding sites manifests as strong intra-motif dependencies between positions 7 and 12, 7 and 11, 5 and 7, and 11 and 12. However, all partitions show a considerable fraction of CpG dinucleotides at positions 9/10, which are methylation sensitive to different degrees. Partition 3 (counted form top) exhibits an additional CpG at positions 11/12 with moderate frequency and methylation sensitivity. While each of these partitions could be modelled decently by its individual (methylation-aware) PWM model, only dependency models as proposed in this study are capable of capturing such highly heterogeneous binding landscapes without prior knowledge about their specific structure.

### Methylation-aware models may explain differential binding

Having established that incorporating methylation-aware genomes and/or intra-motif dependencies is often beneficial for modeling TF binding sites, we further investigate to which extent these models are capable of explaining differential binding across cell types as outlined in Fig. 8A.

**Fig. 8.**
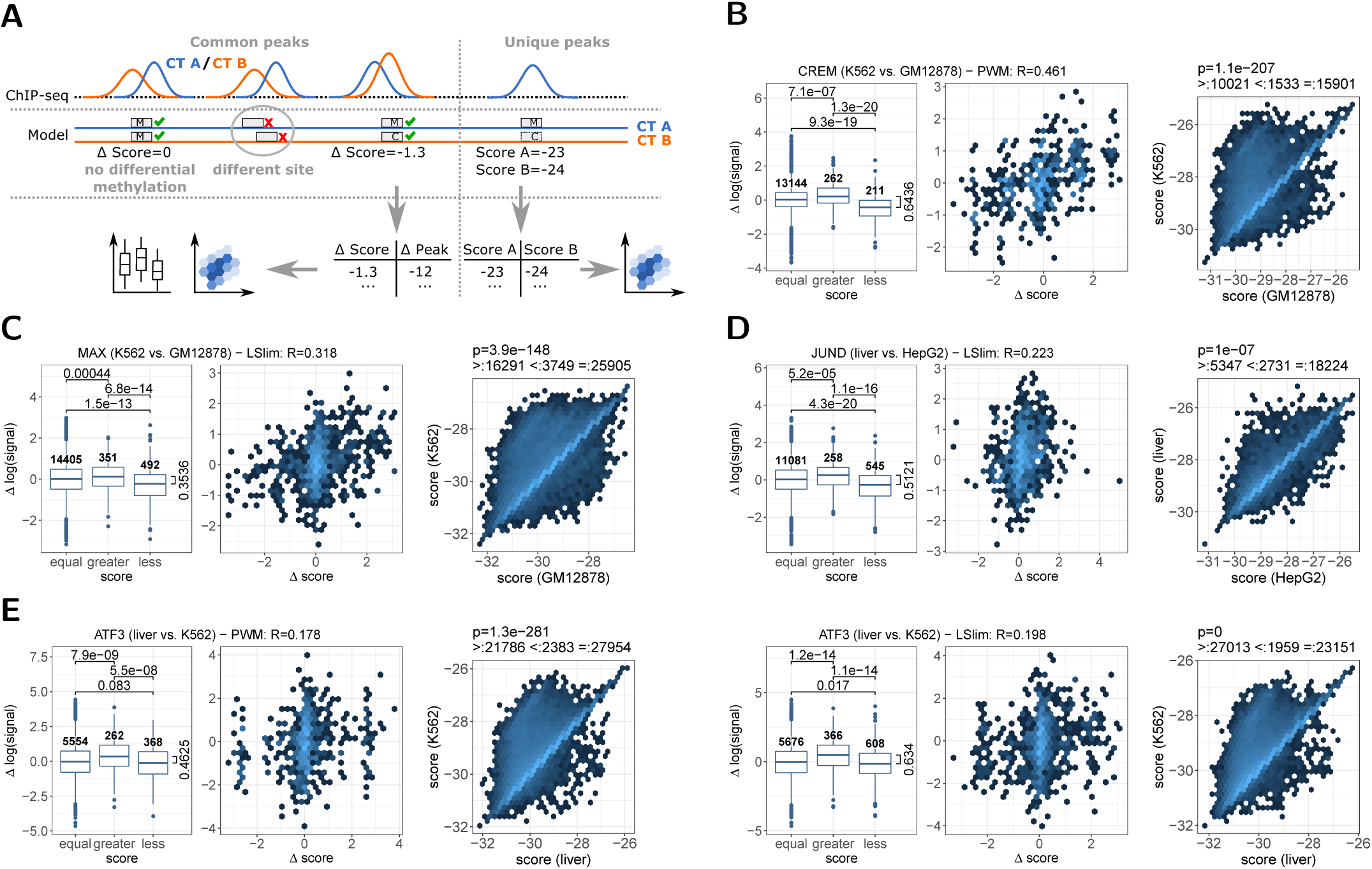
Association of differential model scores and differential binding according to ChIP-seq data. (A) Evaluation schema. For common peaks of two cell types, we consider predicted binding sites at the same location that may show differential methylation and, consequently, different model scores in the methylation-aware genomes. For the peaks containing such binding sites, we record the difference of model scores and the difference in peak height (signal). For unique peaks present in only one of the cell types, we record the scores of the binding sites predicted in the two methylation-aware genomes. (B) Evaluation of cell type-specific binding for CREM in K562 and GM12878 cell types. Left: Comparison of the difference in log signal for binding sites with an equal score in the methylation-aware genomes of K562 and GM12878, with a larger score in K562 than in GM12878, and vice versa. Number of peaks in each group are given above the boxes, p-values from a one-sided Wilcoxon rank sum test are above the boxplots, and the difference of median values between the “greater” and “less” group is indicated. Middle: For those sites with a prediction score differing between K562 and GM12878, we find a correlation of 0.461 between the difference of the log signals and the difference of the prediction scores in those two cell types. Right: Hexbin representation of the scatter plot of scores determined from binding sites in the two methylation-aware genomes for peaks that are present only in K562. Hexbin colours in log scale. (C) Same as (B), but for MAX in K562 and GM12878 cell types. (D) Same as (B), but for JUND in liver and HepG2 cell types. (E) Same as (B), but for ATF3 in liver and K562 cell types using a PWM model (left group) or an LSlim model (right group).

We consider TFs for which ChIP-seq data are available for two cell types. The idea is to compare whether differences in peak occurrence or ChIP-seq signal in both cell lines can be related to a change in binding scores according to our models. In addition to our models, we consider a simple baseline model, which considers average methylation levels of larger genomic regions (cf. Methods) instead of scores of individual binding sites. To associate binding scores with ChIP-seq peaks, we consider the binding sites under ChIP-seq peaks as predicted by the same model, which may have been trained on data from one of the cell types considered or from another cell type. We partition the peaks into “common peaks”, i.e., peaks that are overlapping between the two cell types, and “unique peaks”, i.e., peaks that are present only in one of the cell types.

For the common peaks, and associated binding sites and prediction scores, we separate peaks into those without differential methylation in the binding site and, accordingly, identical prediction scores (“equal”), those with a greater score in cell type A than in cell type B (“greater”), and vice versa (“less”). In addition, we compute the difference in log peak height (“signal”) for each pair of overlapping peaks. If the model could explain differential binding, we would expect these differences to be lower than 0 for the “less” group, around 0 for the “equal” group, and above 0 for the “greater” group, and we test pairwise differences in the distribution of log signal values accordingly by a one-sided Wilcoxon rank sum test.

Boxplots representing this analysis for TF CREM in K562 and GM12878 cell types using a PWM model trained from K562 data (cf. Supplementary Table 3) are shown in the left panel of Fig. 8B. We find significant differences in log signal between all pairs of groups. The difference between the median values for the “less” and “greater” group is 0.6436, which corresponds to a 1.9-fold increase in the ratio of the cell type-specific signal values. Hence, the model appears to be capable of predicting if a peak is larger in cell type A than in cell type B, although the large number of confounding factors, including chromatin accessibility, leads to pronounced variation within each of the groups.

In addition, we plot the differences in log signal against the differences in associated prediction scores and compute the Pearson correlation coefficient between both quantities as shown in the middle panel of Fig. 8B. Here, we exclude peaks without differential methylation in the binding site, since these would obtain a fixed score difference of 0. In case of CREM, we find a substantial correlation between both quantities, although only a small subset of common CREM peaks (473 peaks) participates in the analysis. This may indicate that the model is not only capable of predicting the direction of the change in peak height, but that the difference in prediction scores is associated with the magnitude of this change.

For the unique peaks present only in cell type A, we complement the predicted binding site in the methylation-aware genome of cell type A with the corresponding site in the methylation-aware genome of cell type B, and compute the model scores for both site variants. If DNA methylation as captured by the model could explain the presence and absence of a peak, respectively, we would expect the score for cell type A to be larger than for cell type B. In the right panel of Fig. 8B, we show a hexbin representation of the scatter plot of such pairs of prediction scores for CREM in K562 and GM12878 cell types. Indeed, we find a larger score for K562 than GM12878 for 10021 sites, whereas the opposite is true only for 1533 sites. For the majority of 15901 sites, prediction scores in the methylation-aware genomes of both cell types are identical. Still, the pairwise difference in scores is significantly different from 0 in a Wilcoxon signed rank test (*p* = 1.1 *×* 10^−207^).

In complete analogy, we present results for TF MAX in K562 and GM12878 cell types in Fig. 8C. Here, we consider an LSlim model trained on data for cell type HepG2, i.e., in this case the training cell type is different from the two cell types considered in this analysis. Again, we find significant differences between the three groups of peaks divided by the difference in prediction scores. However, the difference of median values between the “less” and “greater” groups is only 0.3536 in this case. Here, the correlation analysis shows a slightly lower Pearson correlation than for CREM as well with a visible enrichment of score differences around 0. Considering unique peaks, we find approximately 4-fold as many peaks with larger prediction scores in K562 than in GM12878 for peaks that are present only in K562.

Similar tendencies may be observed for JUND in liver and HepG2 cell types using a LSlim model trained from K562 data (Fig. 8D). However, the results for the unique peaks are less pronounced in this case with only 2-fold difference in the number of peaks with greater and lower scores in liver than in HepG2, respectively.

Finally, we illustrate the impact of modelling intra-motif dependencies, i.e., the comparison of PWM and LSlim models, for ATF3 binding sites in liver and K562 cell types in Fig. 8E. While we observe a clear advantage of the LSlim model over the PWM model for all three analyses, this advantage is less pronounced than it had been for the classification-based benchmarks in previous sections.

In Supplementary Figures 4 to 13, we provide results for these and further TFs, and compare these against the baseline model that considers average methylation levels in the regions under the ChIP-seq peaks. It is well known that methylation levels in broader regions, especially in enhancers, are highly informative of TF binding (66). In addition, binding models consider only 20 bp of DNA, which makes the presence of differential methylation less likely than for the simple model. Hence, we expect this to be a strong baseline model. For the common peaks, we indeed find that the differences between the “equal”, “greater” and “less” groups often obtain lower p-values for the baseline than for the methylation-aware binding models, partly due to the larger number of regions with differences in methylation levels. Notably, the binding models often surpass the baseline models for the correlation analysis. Regarding unique peaks, binding models often more clearly show an enrichment of larger scores for the cell type with a peak being present.

In summary, our results suggest that models of TF binding sites learned from methylation-aware genomes and incorporating intra-motif dependencies may indeed be indicative of presence or absence of a ChIP-seq peak and its peak height, despite the many confounding factors that are not related to DNA methylation but strongly influence TF binding.

## CONCLUSIONS

In this paper, we present MeDeMo, a novel framework for TF motif discovery and TFBS prediction that combines information about DNA methylation with models capturing intra-motif dependencies. In contrast to mEpigram (34), MeDeMo does not use a beta value cut-off of 0.5 to obtain a discrete methylation value. Instead, we model the distribution of all beta values using the BETAMIX (40) software to select the cut-off in an informed way. Similar to previous approaches (33, 36), MeDeMo uses an extended 6-letter alphabet with separate symbols for methylated cytosines and the corresponding guanosines on the opposite strand. Also, mEpigram uses a PWM based approach, neglecting intra-motif dependencies. Therefore, MeDeMo using PWM models can be seen as an improved instantiation of mEpigram, but also allows for including intra-motif dependencies when applying LSlim to methylation-aware input data. In agreement with our results, Ngo *et al*. (34) showed that mEpigram outperforms the MEME suite for motif discovery which does not take DNA methylation into account. Previous and this work have shown that many TFs show sensitivity to the status of CpG methylation. Interestingly, it was also shown that enzymes such as DNase1 and the Tn5 transposase, the two most often used enzymes for the measurement of open-chromatin, show differences in DNA cutting or insertion with respect to CpG methylation (7, 67). Thus in genome-wide analysis of such data, neglecting the status of DNA methylation may be harmful in two ways. Binding may be impaired due to TFs that show reduced binding of methylation and abundance of open-chromatin reads may also be affected. Thus, learning of a TF-specific effect of DNA methylation using open-chromatin data only, should carefully integrate both these aspects.

MeDeMo allows the research community to leverage the vast amounts of TF ChIP-seq and DNA methylation datasets available to elucidate the methylation dependence of hundreds of TFs *in vivo*, without the need of performing additional experiments such as Methyl-Spec-seq (33).

Apart from improving TF binding predictions, MeDeMo could also improve the interpretation of methylation QTLs (meQTLs). Methylation QTLs have been reported before to be associated to changes in TF binding, histone modification and gene expression (68). Using MeDeMo, those associations could be understood at more detail, and our analyses regarding differential binding might be a first step towards this goal. Similarly, our tool could provide valuable additional insights into the vast amount of epigenome-wide association studies (EWAS) (69).

Especially in light of upcoming single cell applications as single-cell methylation (70) and single-cell chromatin accessibility assays become available (71), the need of methylation aware TFBS prediction approaches will rise even further in the near future. MeDeMo will help to fulfill these data analysis needs.

## METHODS

### Data

We downloaded whole genome bisulfite sequencing data for 3 cell-lines (K562, HepG2, GM12878) from ENCODE as well as for 2 replicates of primary human hepatocytes (DEEP). The ENCODE data has been processed following the uniform ENCODE-Processing pipeline, the DEEP data has been processed following the DEEP *MCSv3* pipeline (72). Furthermore, we downloaded TF-ChIP seq peak calls (IDR thresholded peaks) from ENCODE for 336 experiments in K562, 145 in HepG2, 129 in GM12878 and 25 in primary human hepatocytes (liver). Data accession IDs are provided in Supplementary Table 1.

### Generation of methylation-aware genomes

To generate a methylation-aware genome sequence, where a methylated C is replaced by ‘M’ and a G opposite of a methylated C is replaced by ‘H’, we discretized the methylation calls from whole genome bisulfite data using BETAMIX (40) and the parameter *–components unimodal unimodal*.

### Training procedure

Motif models are learned from ChIP-seq data by the discriminative maximum supervised posterior principle within the SlimDimont framework (8, 73). To this end, we use as positive training sets genomic regions under all ChIP-seq positive peaks (optimal IDR thresholded peaks) as downloaded from the ENCODE project and extract the sequence of length 1, 000bp around the peak center. In addition, we use two different sets of negative training sets. First, we randomly draw 10, 000 regions uniformly from the complete genome excluding any ChIP-seq positive region of the TFs studied (random) and again extract the sequence of length 1, 000bp around the center of each region. Second, we consider dinucleotide shuffled versions of each positive sequence in the training set (shuffled). Negative training sequences are weighted such that their total weight equals the number of positive training sequences. In either case, we extract sequences from the original *hg38* genome with standard DNA nucleotides and, alternatively, sequences from the genomes including methylation calls (Section 4.2). As the methylated genomes are cell type-specific we always use those matching the cell type of the corresponding ChIP-seq experiment. Sequences from the negative sets are also extracted from the matching genome versions. Models that are discovered *de novo* from these data sets are i) standard position weight matrices and ii) LSlim models (8) with a maximum distance of 5bp between putatively dependent positions. In general, motif discovery within the SlimDimont framework (8, 73) may report multiple motifs per input data set. For the remainder of the analyses described here, we only consider the first reported motif according to the ranking by the value of the maximum supervised posterior objective function used internally in the SlimDimont framework as proposed previously (73).

### Prediction procedure

Given a trained motif model and an input set of sequences, we compute for each sequence the log-likelihood of all overlapping sub-sequences on both strands matching the motif length. We then chose as predicted value for that sequence the maximum over all these log-likelihood values. In contrast to alternative scores, like the *sum occupancy score* (74) integrating over all log-likelihood values, this procedure makes sure that the score of a sequence can be attributed to one specific sub-sequence with its methylation pattern.

### Cross validation procedure

For benchmarking the different models learned from sequence with and without methylation information, we follow a 10-fold cross validation procedure. Specifically, we partition ChIP-seq positive regions and (for the first training variant) drawn negative regions into 10 equally sized sets, where in each cross validation fold, the union of 9 of these sets is used for training and the remaining set is used for testing. Since partitioning is performed before extracting sequences, training sets in the different cross validation folds are identical (aside from methylation information) between the different genome versions.

### Evaluating performance

For evaluating performance of a model trained on and applied to sequences from a specific genome version, we consider a classification problem discriminating ChIP-seq positive from negative sequences. The positive set comprises all sequences extracted under ChIP-seq positive regions from the corresponding test partition. The negative set, in turn, comprises sequences from genomic regions that are again randomly drawn uniformly from the complete genome, in this case excluding all ChIP-seq positive regions for all TFs studied and also excluding the negative regions used for training. In total, this negative set contains 100, 000 regions, which are again partitioned into 10 test sets to capture variability among different choices of negatives. Given a model, scores for all sequences in positive and negative sets are computed as described in Section 4.4. The ability of these scores to distinguish positives from negatives is then evaluated by the area under the precision recall curve (AUC-PR) as determined by the PRROC R package (43). Models trained on the training partition of one ENCODE data set for one specific TF are evaluated i) on the test partition of the same data set, ii) on the corresponding test partition of other data sets for the same TF and cell type, and iii) on the corresponding test partition of other data sets for the same TF in other cell types. We refer to the first two cases as *within cell type*, and to the latter case as *across cell type*.

### Model visualization

Since parameters of models learned by discriminative learning principles may be skewed to optimize prediction accuracy, a direct visualization of these parameters may lead to un-intuitive results. Hence, we follow the approach of (8) and visualize models based on their predicted binding sites on the training data represented by traditional sequence logos and dependency logos generated by the DepLogo R package (64).

### Methylation sensitivity

We investigate the methylation sensitivity of a trained model again based on predicted binding sites. To this end, we consider models learned from sequences using the extended, methylation-aware alphabet and binding sites predicted from the corresponding training data set. Each of these binding sites is first converted to the standard DNA alphabet replacing occurrences of M with C and of H with G. We use this modified sequence to compute a *base score* without methylation. We then consider each CpG dinucleotide within the sequence (regardless if it was methylated in the original sequence) and change both the C to M and the G to H (MH). We compute the score of the modified sequence according to the model and determine its difference relative to the base score. If this score is larger than the base score, we consider the influence of methylation on such a nucleotide (in this sequence context) as *beneficial*, and as *detrimental* otherwise. For each binding site position, we also compute the relative abundance of CpGs in the predicted binding sites and the average of the score differences of the MH case relative to the base score.

### Differential binding

For analyzing the association between scores of predicted binding sites and differential binding, we consider pairs of cell types, A and B, with ChIP-seq data available for the same TF. In this analysis, we distinguish common peaks that overlap between the two cell types, and unique peaks present in only one of the cell types. We further predict one binding site per ChIP-seq peak at the position yielding the maximum score as described in Section 4.4.

For the common peaks, we only compare (scores of) binding sites that are predicted at exactly the same genomic location in the methylation-aware genomes of both cell types, as this allows for a direct comparison of prediction scores. This requirement is reasonable as, in principle, the position of a predicted binding site could change due to differences in the methylation states of the two cell types. Since only such common binding sites are considered, we identify common peaks by the presence of predictions at identical genomic locations within the two methylation-aware genomes. Predicted binding sites in both cell types are recorded together with the corresponding prediction scores and the peak heights (column 7 of the narrowPeak format) of the surrounding peaks.

We further identify unique peaks for cell type A using the bedtools (75) command “bedtools intersect -v -a peaksA.bed -b peaksB.bed > onlyA.bed”. For binding sites predicted from the methylation-aware genomes of cell type A, we extract the corresponding sequence from the methylation-aware genome of cell type B, and record prediction scores for these two predicted sites. We proceed in complete analogy to identify unique peaks for cell type B.

For these analyses, we aim at using the same models that have also been considered for classification-based benchmarks. However, as these benchmarks are based on 10-fold cross validation experiments, we also obtain a set of 10 models per TF and training data set. For this reason, we perform the above-mentioned procedure for each of the 10 models, and average prediction scores per ChIP-seq peak before proceeding with statistical analysis and visualization.

As a reference, we also consider a simple baseline, which measures methylation levels of the sequences under ChIP-seq peaks. Specifically, we extract sequences of length 1000 bp and determine, on either strand of the DNA sequence, the fraction of cytosines that are methylated according to the methylation-aware genome of a cell type. We center the extracted sequences at the position of the predicted target site instead of the (cell type-specific) peak center or peak summit to ensure that methylation levels in different cell types are measured for the same genomic region.

### Method implementation

We implement the model, training procedure, and prediction procedure based on the existing implementation of the SlimDimont approach (8). The basic modification compared with the version published previously is the extension of the alphabet to A, C, G, T, M and H, where M is complementary to H. This extension allows us to include information about methylation while preserving the possibility to compute reverse complements of input sequences, which is necessary because in ChIP-seq data binding sites may be located on either DNA strand. We provide this methylation-aware toolbox termed MeDeMo for motif discovery as i) stand-alone binary versions with graphical user interface and command line interface (cf. “Availability of data and materials”).

## DECLARATIONS

### Ethics approval and consent to participate

Not applicable.

### Consent for publication

Not applicable.

### Availability of data and materials

All ChIP-seq data sets analyzed in this study are available from ENCODE. ENCODE IDs of the corresponding narrowPeak files are listed in Supplementary Table 1 and can be accessed via the URL schema https://www.encodeproject.org/search/?searchTerm=<ID>. The methylation-aware genome variants and the models generated during the current study are available as a Zenodo archive from https://doi.org/10.5281/zenodo.3723985. The MeDeMo software is available as stand-alone binary versions with graphical user interface and command line interface at http://www.jstacs.de/index.php/MeDeMo. The source code of the software is available from github at https://github.com/Jstacs/Jstacs. Classes specifically implemented for this project are provided in package `projects.methyl`.

### Authors’ contributions

FS and MHS originally conceived the idea of the study. JG, FS and MHS designed the experiments. FS and JG designed and implemented the MeDeMo workflow. FS and JG prepared the figures. All authors contributed to paper writing.

## Acknowledgements

We thank Christopher Schröder for help to run BETAMIX. We thank Ekaterina Shelest for valuable discussions. We thank Yang Gao for help and discussions in an initial test study.

This preprint is formatted using a LATEX class by Ricardo Henriques.

