## Supplementary Tables and Figures for "Widespread effects of DNA methylation and intra-motif dependencies revealed by novel transcription factor binding models"

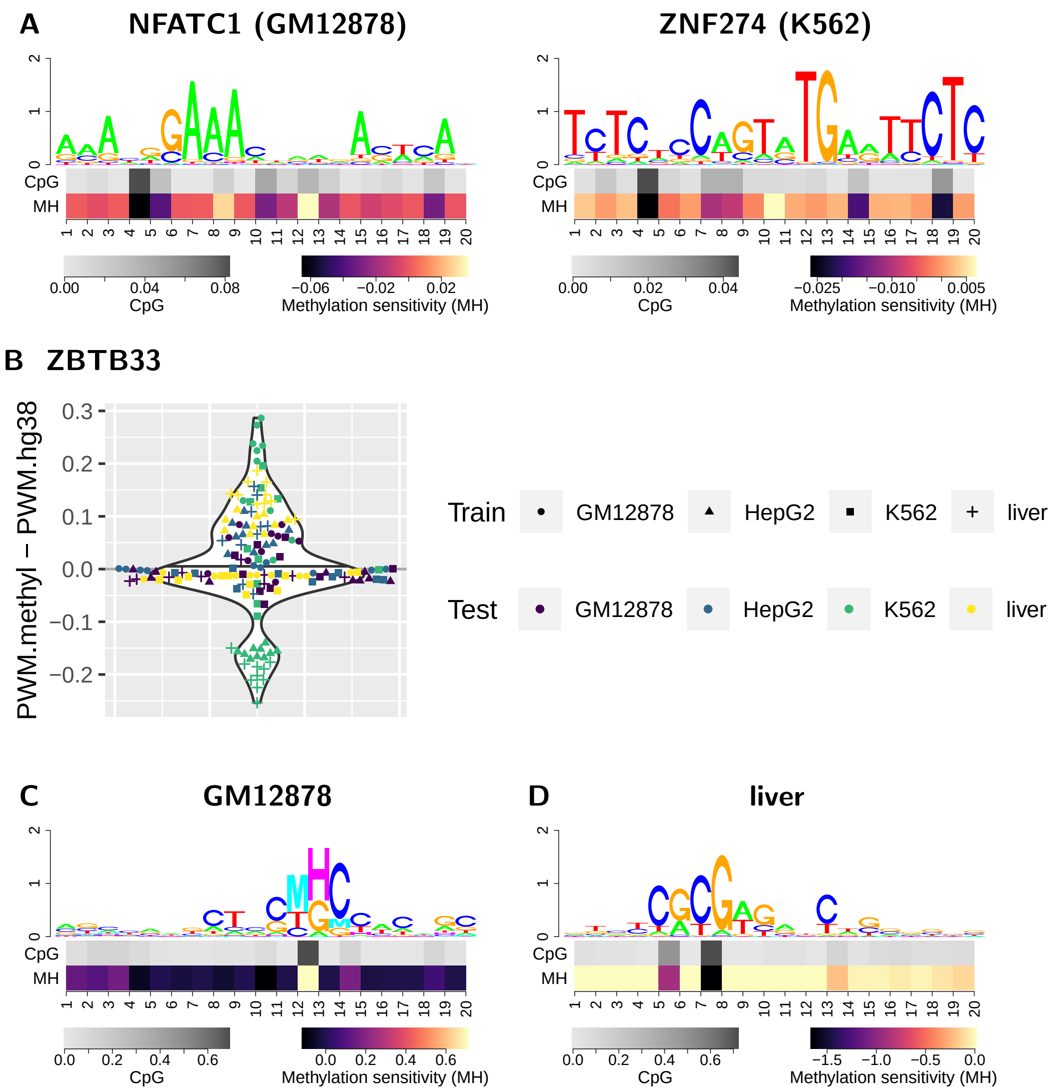


Figure 1: Methylation sensitivity is positive for TFs with known preference for methylated DNA. (A) For NFATC1, we find methylation sensitivity values that are clearly positive at some positions (8,12), but also clearly negative at other positions, while the amplitude of methylation sensitivity and CpG content is rather low. Similarly, we find methylation sensitivity values that are clearly positive at some positions (especially position 10), but also clearly negative at other positions for ZNF274. (B) For ZBTB33, we include within cell type and across cell type comparisons in a single plot, which compares the performance of a methylation-aware PWM model with the corresponding PWM model learned on the original hg38 genome. For within cell type performance, we find an improvement when including methylation information for GM12878 (dark violet dots), HepG2 (blue triangles), and liver (yellow crosses), whereas results for K562 (green squares) are rather mixed. Across cell types , the motif trained on methylation-aware GM12878 data works particularly well for K562 (green dots), but less well for HepG2 (blue dots) and liver (yellow dots) data. In turn, the motif trained on methylation-aware liver data works well for HepG2 (blue crosses) but not for GM12787 (dark violet crosses) and K562 (green crosses) data. (C) Non-canonical motif discovered for ZBTB33 in GM12878 cells with clear methylation preference, which also performed well on K562 data. For K562, a strong methylation preference for ZBTB33 has been reported before (1). (D) The motif discovered from liver data resembles the canonical ZBTB33 motif, but shows a negative effect of binding site methylation. This motif also worked well for HepG2 data.

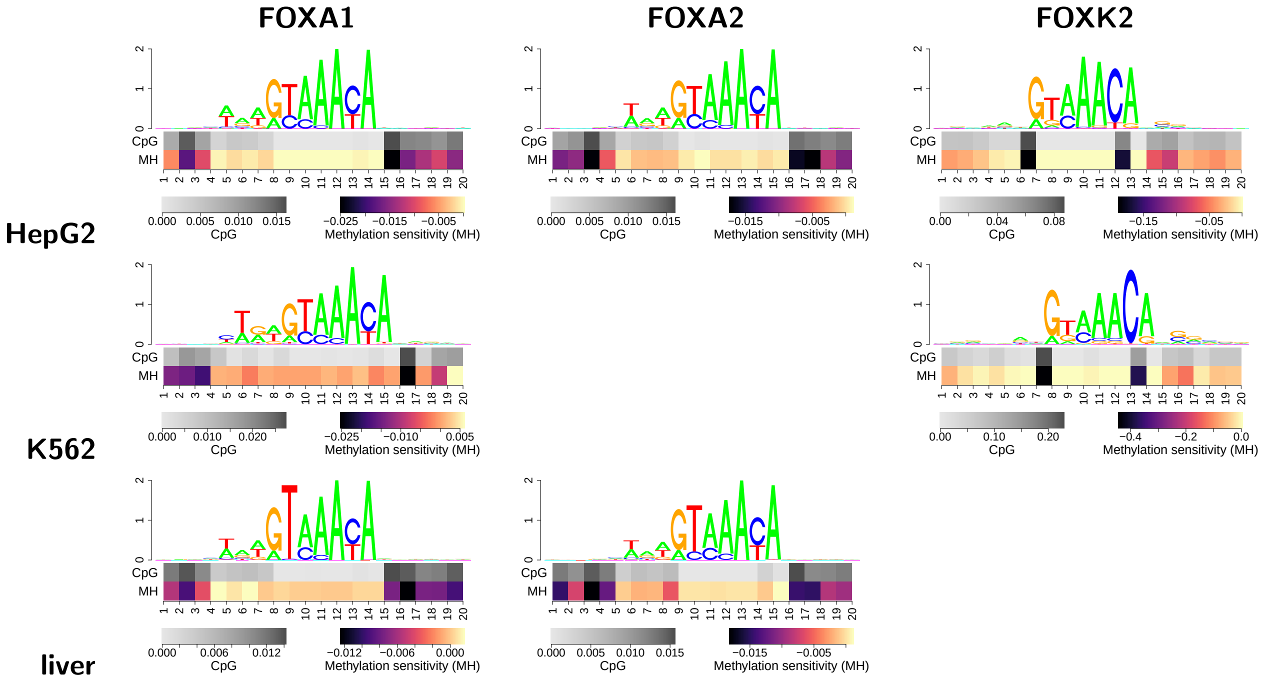


Figure 2: Methylation sensitivity may differ between members of a TF family. While methylation sensitivity of FOXA1 and FOXA2 is highly similar in all cell types, that of FOXK2 is noticeably different, although the motifs of all three TFs appear to be highly similar.

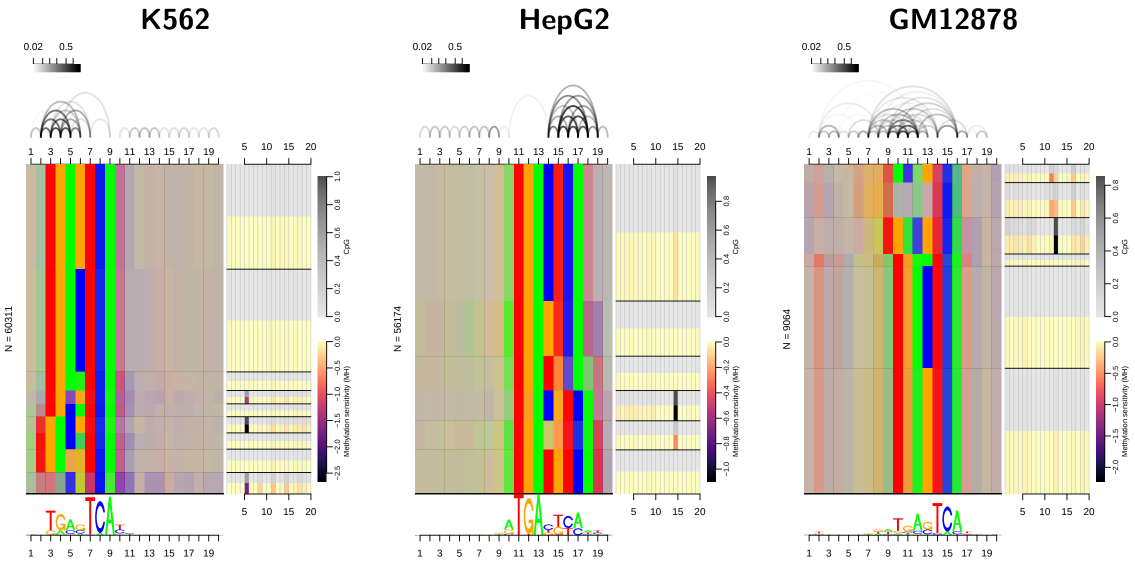


Figure 3: Dependency logos and profiles of methylation sensitivity for JUND in three cell types (K562, HepG2, GM12878). In all three cases, we consistently find both, the long-spacer and the short-spacer variant, of the binding motif, with a specific pattern of methylation sensitivity, which is prominent only for the long-spacer variant.

Table 1: List of the Encode IDs of all ChIP-seq data sets used in this study for the four cell types GM12878, HepG2, K562, and liver.

| TF | GM12878 | K562 | HepG2 | liver |
|---|---|---|---|---|
| AFF1 | | ENCFF489SKQ, ENCFF869BYK | | |
| AGO1 | | ENCFF100VYA | ENCFF627BHP | |
| AGO2 | | | ENCFF465FII | |
| ARHGAP35 | | ENCFF089PKE | | |
| ARID1B | | ENCFF249TYS | | |
| ARID2 | | ENCFF344MKI | | |
| ARID3A | ENCFF003VDB | ENCFF757OML | ENCFF247GXE | |
| ARNT | ENCFF758RQJ | ENCFF447FIO, ENCFF655EFA, ENCFF913AQF | ENCFF616WXJ | |
| ASH1L | | ENCFF958YSG | | |
| ASH2L | ENCFF096XRG | | ENCFF638IUM | |
| ATF2 | ENCFF210HTZ | ENCFF803FHN | ENCFF089BQU | |
| ATF3 | | ENCFF467WOR, ENCFF937OKC | ENCFF137OEY | ENCFF146URA, ENCFF782SGI |
| ATF4 | | ENCFF182MNO | | |
| ATF7 | ENCFF495PWL | ENCFF371SJR | ENCFF498YGH | |
| ATM | | | ENCFF906FVB | |
| BACH1 | ENCFF725YZH | | | |
| BATF | ENCFF832YIE | | | |
| BCL11A | ENCFF383HAY | | | |
| BCL3 | ENCFF247MHT | | | |
| BCLAF1 | ENCFF587BJK | ENCFF054DTJ, ENCFF094XPM, ENCFF496YJC | ENCFF506PXL | |
| BCOR | | ENCFF186JKG | | |
| BHLHE40 | ENCFF370ZNL | ENCFF477JTV | ENCFF361YXC, ENCFF863ATX | |
| BMI1 | ENCFF592LPO | ENCFF352DRR | | |
| BRCA1 | ENCFF005JKU | ENCFF652NES | ENCFF897ETK | |
| BRD4 | | ENCFF806CQB | ENCFF736GHL | |
| BRD9 | | ENCFF411RMT | | |
| CBFA2T2 | | ENCFF419PEK | | |
| CBFA2T3 | | ENCFF153IFH | | |
| CBFB | ENCFF070SOX | | | |
| CBX1 | | ENCFF163FLA | | |
| CBX2 | | | ENCFF501QII | |
| CBX3 | ENCFF552QOA | ENCFF951BQB | | |
| CBX5 | ENCFF417SVR | ENCFF403TAE | | |
| CC2D1A | | ENCFF180TUM | | |

Table 1: (continued)

| TF | GM12878 | K562 | HepG2 | liver |
|---|---|---|---|---|
| CCAR2 | | ENCFF704PGT | ENCFF039LHY | |
| CDC5L | | ENCFF384ALH | | |
| CEBPB | ENCFF786YYI | ENCFF321KQD | ENCFF862DXR, ENCFF915ZYE | |
| CEBPBZ | | ENCFF797OWK | | |
| CEBPZ | ENCFF243GOG | | ENCFF195BKI | |
| CHAMP1 | | ENCFF646MEF, ENCFF919KNQ | | |
| CHD1 | ENCFF863CTN | | | |
| CHD2 | ENCFF546AYN | | ENCFF181XMM | |
| CHD4 | ENCFF249SIN | | ENCFF148ABR | |
| COPS2 | | ENCFF552EBC | | |
| CREB1 | | | ENCFF550TXR | |
| CREB3L1 | | ENCFF566HGU | | |
| CREM | ENCFF091YID | ENCFF021XJN | ENCFF290UGF | |
| CTBP1 | | ENCFF349UTF | | |
| CTCF | ENCFF356LIU | ENCFF119XFJ, ENCFF396BZQ, ENCFF519CXF, ENCFF843VHC | ENCFF543WTP | |
| CUX1 | ENCFF567NFS | ENCFF556HMX | | |
| DACH1 | | ENCFF870LJV | | |
| DDX20 | | ENCFF536LKB | | |
| DEAF1 | | ENCFF532HCE | | |
| DNMT1 | | ENCFF549TVW | | |
| DPF2 | ENCFF771IAW | ENCFF217ZTP, ENCFF537VKZ | | |
| E2F1 | | ENCFF134JLR, ENCFF445VTT | | |
| E2F4 | ENCFF687SFB | | | |
| E2F6 | | ENCFF533GSH | | |
| E2F7 | | ENCFF013EHI | | |
| E2F8 | ENCFF412GFI | ENCFF171WWF | | |
| E4F1 | ENCFF035GFS | ENCFF752KNU | | |
| EBF1 | ENCFF249SVT | | | |
| EED | ENCFF023ALY | | | |
| EGR1 | | ENCFF175VSS, ENCFF375RDB, ENCFF561OGS | | ENCFF617JQS, ENCFF808WST |
| EHMT2 | | ENCFF682XPD | ENCFF413RQL | |
| EKL1 | ENCFF432AQP | | | |
| ELF1 | ENCFF948CPI | ENCFF617ZLL | ENCFF840RWO | |

Table 1: (continued)

| **TF** | **GM12878** | **K562** | **HepG2** | **liver** |
|---|---|---|---|---|
| ELF4 | | ENCFF539SXG | | |
| ELK1 | | ENCFF119SCQ | | |
| EP300 | ENCFF510FUM | ENCFF755HCK | ENCFF674QCU, ENCFF806JJS | |
| EP400 | | ENCFF225BXA | | |
| ESRRA | ENCFF722LJP | ENCFF592GWM | | |
| ETS1 | ENCFF980VOD | ENCFF461PRP | ENCFF128TUP | |
| ETV4 | | | ENCFF710CRT | |
| ETV6 | ENCFF116AMK | ENCFF426GSY, ENCFF658SGJ | | |
| EWSR1 | | ENCFF560CYG | | |
| EZH2 | ENCFF615NYO | | ENCFF504QZJ | |
| FIP1L1 | | ENCFF084DTV | ENCFF031LBW | |
| FOSL1 | | ENCFF087MFG | | |
| FOSL2 | | | ENCFF054ESU | |
| FOXA1 | | ENCFF765NAN | ENCFF152BOT, ENCFF367TQC, ENCFF872MGU | ENCFF324QGE, ENCFF951VPZ |
| FOXA2 | | | ENCFF184NAC, ENCFF259BJR | ENCFF168JLI, ENCFF293LRQ |
| FOXK2 | ENCFF990MTR | ENCFF490EQR | ENCFF315CHX | |
| FOXM1 | | ENCFF778PWE | | |
| FOXP1 | | | ENCFF029UJC | |
| FUS | | ENCFF688ARM | ENCFF216YZI | |
| GABPA | ENCFF946ACA | ENCFF124HAC | ENCFF054HJA | ENCFF280YAF, ENCFF344XWK |
| GABPB1 | | ENCFF700DXR | | |
| GATA1 | | ENCFF148JKK | | |
| GATA2 | | ENCFF173TXA | | |
| GATA4 | | | ENCFF097OXR | |
| GATAD2A | | ENCFF950ZWP | | |
| GATAD2B | ENCFF298AIX | ENCFF569CMJ | | |
| GMEB1 | | ENCFF678VPQ | | |
| GTF2F1 | | ENCFF478HYJ, ENCFF713ALB, ENCFF843UHP | ENCFF599TWF, ENCFF876GXQ | |
| HCFC1 | ENCFF722QBB | ENCFF167RXK | ENCFF485SRU | |
| HDAC1 | | ENCFF188TBM, ENCFF557WXK, ENCFF661VOO, ENCFF758PGF | ENCFF069KPS | |

Table 1: (continued)

| TF | GM12878 | K562 | HepG2 | liver |
|---|---|---|---|---|
| HDAC2 | ENCFF299UPZ | ENCFF363GSV, ENCFF519RWJ, ENCFF618YRQ | ENCFF182XZZ, ENCFF589GSN | |
| HDAC3 | | ENCFF742LSD | | |
| HDAC6 | | ENCFF295GBP | ENCFF109EXK | |
| HDGF | ENCFF442WRJ | ENCFF297WMY, ENCFF575WFB | | |
| HES1 | | ENCFF010OOE | | |
| HMBOX1 | | ENCFF718DFX | | |
| HNF1A | | | ENCFF800QTO | |
| HNF4A | | | ENCFF072CXB | ENCFF837QHJ, ENCFF905JAC |
| HNF4G | | | ENCFF086CTA | ENCFF497MUF |
| HNRNPH1 | | ENCFF844QFF | ENCFF046NUR | |
| HNRNPK | | ENCFF984QUV | ENCFF828KXG | |
| HNRNPL | | ENCFF984ESZ | ENCFF039CUI | |
| HNRNPLL | | ENCFF662WPN | ENCFF890KTX | |
| HNRNPUL1 | | ENCFF991ZSC | ENCFF509YFF | |
| HSF1 | ENCFF603BID | | | |
| IKZF1 | ENCFF968NOG | ENCFF785BTP, ENCFF994OQH | ENCFF969BZA | |
| IKZF2 | ENCFF088OLI | | | |
| ILF3 | | ENCFF368AAQ | | |
| IRF2 | | ENCFF886EVL | | |
| IRF3 | ENCFF604AZX | | | |
| IRF4 | ENCFF720YMW | | | |
| IRF5 | ENCFF843HDK | | | |
| JUN | | ENCFF394CEC | | |
| JUNB | ENCFF478XNA | ENCFF739XTO | | |
| JUND | ENCFF873DJD | ENCFF213EYD | ENCFF430PEI, ENCFF539GRW | ENCFF229COM |
| KAT2A | ENCFF710ROZ | | | |
| KAT2B | | | ENCFF091BEK | |
| KAT8 | | ENCFF207ZEK | | |
| KDM1A | ENCFF799KZP | ENCFF483BRD, ENCFF728KKP, ENCFF796VMI | ENCFF768FGG | |
| KDM4B | | ENCFF470RHZ, ENCFF955AOD | | |
| KDM5A | | | ENCFF334HKG | |
| KDM5B | | ENCFF668XLN | | |
| KLF16 | | ENCFF379LKE | | |

Table 1: (continued)

| TF | GM12878 | K562 | HepG2 | liver |
|---|---|---|---|---|
| KLF5 | ENCFF417WPC | | | |
| L3MBTL2 | | ENCFF423LPW | | |
| LCORL | | | ENCFF611PIO | |
| LEF1 | | ENCFF134HQP, ENCFF697VRJ | | |
| MAFF | | ENCFF498MGH | ENCFF493TIR | |
| MAFK | ENCFF186AWV | ENCFF893SCL | ENCFF171OJF, ENCFF770TZL | |
| MAX | ENCFF270NAL | ENCFF618VMC, ENCFF900NVQ | ENCFF140PUO | ENCFF669BQN |
| MAZ | ENCFF348STZ | | ENCFF144TBQ | |
| MBD2 | | ENCFF617QSK | | |
| MCM2 | | ENCFF043HHG, ENCFF571REC | | |
| MCM3 | | ENCFF672PYP | | |
| MCM5 | | ENCFF603SXI, ENCFF658SJY | | |
| MCM7 | | ENCFF159MQI, ENCFF288ZRD, ENCFF914ELA | | |
| MEF2A | ENCFF958GXF | ENCFF310SMW | | |
| MEF2B | ENCFF623FAW | | | |
| MEF2C | ENCFF830BRO | | | |
| MEIS2 | | ENCFF937UEE | | |
| MGA | | ENCFF525MPI | | |
| MIER1 | | ENCFF163YZB | | |
| MITF | | ENCFF071NYD, ENCFF262TMM | | |
| MLLT1 | ENCFF125MEN | ENCFF010AIG, ENCFF388LUX | | |
| MNT | | ENCFF454QQD, ENCFF459DYU, ENCFF926CRV | ENCFF482JSR, ENCFF562FMQ | |
| MTA1 | | ENCFF801KEW | | |
| MTA2 | ENCFF587POH | ENCFF558XIL, ENCFF713ZVD | | |
| MTA3 | ENCFF661FMB | ENCFF459XLR | | |
| MXI1 | ENCFF199HGX | ENCFF243QTL | | |
| MYB | ENCFF402TSJ | | | |
| MYBL2 | | ENCFF905KOD | | |
| MYC | | ENCFF492XUU | | |
| MYNN | | ENCFF272LLG | | |
| NBN | ENCFF811VEN | | ENCFF516UWH | |

Table 1: (continued)

| TF | GM12878 | K562 | HepG2 | liver |
|---|---|---|---|---|
| NCOA1 | | ENCFF382RFJ, ENCFF474QDS, ENCFF589OOF | | |
| NCOA2 | | ENCFF071SOH, ENCFF584SNZ | | |
| NCOA4 | | ENCFF749HKV | | |
| NCOA6 | | ENCFF438BWN | | |
| NCOR1 | | ENCFF856HUK | ENCFF616RSZ | |
| NEUROD1 | | ENCFF755APC | | |
| NFATC1 | ENCFF138ZBJ | | | |
| NFATC3 | ENCFF704PDA | ENCFF082EPO, ENCFF430JFH | | |
| NFE2 | | ENCFF312XHI | | |
| NFE2L2 | | | ENCFF882YLO | |
| NFIC | ENCFF480WDX | ENCFF092TVM | | |
| NFKB | | | ENCFF162TPR | |
| NFRKB | | ENCFF158FUG, ENCFF779KIS | | |
| NFXL1 | ENCFF860IXB | ENCFF329STX | | |
| NFYA | ENCFF278GJK | | | |
| NFYB | ENCFF510NDO | | | |
| NONO | | ENCFF515YFU, ENCFF823CQK | ENCFF108IZQ, ENCFF420QKI | |
| NR0B1 | | ENCFF305OOU | | |
| NR2C1 | ENCFF462AKP | ENCFF023XHV | | |
| NR2C2 | ENCFF434HVY | ENCFF791ZPU | | |
| NR2F1 | ENCFF531KOV | ENCFF363IQN | | |
| NR2F2 | | ENCFF118HUH | | ENCFF379TVQ |
| NR2F6 | | ENCFF194VBK | ENCFF350CKI | |
| NR3C1 | | ENCFF315MUH, ENCFF821YMC | | |
| NRF1 | ENCFF652BRY | ENCFF543STN, ENCFF626VDA, ENCFF782YFS | ENCFF313RFR, ENCFF418DKQ | |
| NUFIP1 | | ENCFF885JMZ | | |
| PAX5 | ENCFF196JGP | | | |
| PAX8 | ENCFF992JWY | | | |
| PBX3 | ENCFF926LHG | | | |
| PCBP1 | | ENCFF467RYH | ENCFF487WAN | |
| PCBP2 | | ENCFF941XZW | ENCFF642XRH | |
| PHB2 | | ENCFF988OXX | ENCFF882RPA | |
| PHF20 | | ENCFF259HUS | | |

Table 1: (continued)

| TF | GM12878 | K562 | HepG2 | liver |
|---|---|---|---|---|
| PHF21A | | ENCFF657UVA | | |
| PHF8 | | ENCFF952YDR | ENCFF202WIO | |
| PKNOX1 | ENCFF335ADU | ENCFF062VBB | | |
| PLRG1 | | | ENCFF873OHG | |
| PML | | ENCFF800QDU | | |
| POU5F1 | | ENCFF814QPF | | |
| PRDM10 | | ENCFF600HPZ | | |
| PRPF4 | | ENCFF417RQZ | ENCFF908QCS | |
| PTBP1 | | ENCFF917HXV | ENCFF875ZPV | |
| PYGO2 | | ENCFF442XXV | | |
| RAD21 | ENCFF654EGO | | ENCFF093XOJ, ENCFF874VFZ | ENCFF229WFR |
| RAD51 | ENCFF996NBR | ENCFF740OPF | ENCFF859MBC | |
| RB1 | ENCFF034OSV | ENCFF328QZM | | |
| RBBP5 | ENCFF687SSY | ENCFF666PCE | | |
| RBFOX2 | | ENCFF232ASB | ENCFF871YRG | |
| RBM14 | | ENCFF465UMU | | |
| RBM15 | | ENCFF563WDZ | | |
| RBM17 | | ENCFF056OIG | | |
| RBM22 | | ENCFF420IBN | ENCFF305WYD | |
| RBM25 | | ENCFF102XVH | | |
| RBM34 | | ENCFF670ILH | | |
| RBM39 | | ENCFF503DIK | ENCFF420ALF | |
| RCOR1 | ENCFF470ZMK | ENCFF968SUH | ENCFF987VKU | |
| RELB | ENCFF105YDI | | | |
| REST | ENCFF313CII | ENCFF023ZUW, ENCFF290ESJ | ENCFF669XCW, ENCFF986RRJ | ENCFF178WRO, ENCFF288XHG |
| RFX1 | | ENCFF193PVX, ENCFF905GXS | ENCFF788CJF | |
| RFX5 | ENCFF259LNG | ENCFF201YKU | ENCFF059GWW | |
| RLF | | ENCFF599CBB | | |
| RNF2 | | ENCFF349MSP, ENCFF462AZY, ENCFF741CLJ, ENCFF820LKT | ENCFF380SYL | |
| RUNX1 | | ENCFF091MQJ, ENCFF545WXN | | |
| RUNX3 | ENCFF677QUK | | | |
| RXRA | ENCFF313BDA | | ENCFF105TFM | ENCFF201KGJ |
| SAFB | | ENCFF411YVY | | |
| SAFB2 | | ENCFF087DKT | | |
| SAP30 | | ENCFF103RHL | | |

Table 1: (continued)

| TF | GM12878 | K562 | HepG2 | liver |
|---|---|---|---|---|
| SETDB1 | | ENCFF690WNQ | | |
| SIN3A | ENCFF050CYK | ENCFF407VGB, ENCFF802JAN | ENCFF635YMI | |
| SIN3B | | ENCFF543INR | ENCFF193DQZ | |
| SIX5 | ENCFF864TFH | ENCFF247LOF | | |
| SKI | | | ENCFF035ZFO | |
| SKIL | ENCFF903KEI | ENCFF254QDM | | |
| SMAD1 | ENCFF987PGY | ENCFF084BUP | | |
| SMAD2 | | ENCFF186MFI | | |
| SMAD5 | ENCFF855SJG | ENCFF069AAY | | |
| SMARCA4 | | ENCFF361RWX, ENCFF703NAE, ENCFF868UOJ | | |
| SMARCA5 | ENCFF052STI | ENCFF481TNF | | |
| SMARCB1 | | ENCFF308QHX | | |
| SMARCC2 | | ENCFF751ZVX | ENCFF150NHK | |
| SMARCE1 | | ENCFF435SZS | ENCFF210HAA | |
| SMC3 | ENCFF572RPI | ENCFF175UEE | ENCFF035YWE | |
| SNIP1 | | ENCFF529BDW | | |
| SNRNP70 | | ENCFF206MJS | ENCFF858FBZ | |
| SOX13 | | | ENCFF257QND | |
| SOX6 | | ENCFF431STY | ENCFF944LNI | |
| SP1 | | ENCFF452LDK | ENCFF175VXL, ENCFF735WMX | ENCFF433EFF, ENCFF978TMH |
| SPI1 | ENCFF071ZMW | ENCFF414ECK | | |
| SREBF1 | | ENCFF777MYW | | |
| SRF | ENCFF182IFE | | | |
| SRSF3 | | ENCFF926XGK | | |
| SRSF4 | | | ENCFF122FVR | |
| SRSF7 | | ENCFF550VUN | | |
| SRSF9 | | ENCFF217HAW | ENCFF121PED | |
| STAT1 | ENCFF323QQU | | | |
| STAT3 | ENCFF923CHO | | | |
| STAT5A | | ENCFF517IXK | | |
| SUPT20H | ENCFF069YVD | | | |
| SUPT5H | | ENCFF721DPQ | | |
| SUZ12 | | ENCFF440XZI, ENCFF856HYC | ENCFF239LRW | |
| SYNCRIP | | | ENCFF157IIV | |
| TAF1 | ENCFF540AAP | | ENCFF234TBW | ENCFF214OJW |
| TAF15 | | ENCFF710LLF | ENCFF718RXL | |
| TAF7 | | ENCFF852NOL | | |

Table 1: (continued)

| TF | GM12878 | K562 | HepG2 | liver |
|---|---|---|---|---|
| TAF9B | | ENCFF223HDM | | |
| TAL1 | | ENCFF078OUD, ENCFF475LFH | | |
| TARDBP | ENCFF668JHK | ENCFF448YOS, ENCFF641AXD, ENCFF909RMQ | ENCFF696QPP | |
| TBL1XR1 | ENCFF392JWA | ENCFF239WFN, ENCFF868SWL | ENCFF126KGW | |
| TBP | ENCFF896UZB | ENCFF370YGS | ENCFF534GKQ | |
| TBX21 | ENCFF971VHK | | | |
| TBX3 | | | ENCFF654KVO, ENCFF887DUY | |
| TCF12 | ENCFF768VSH | ENCFF912LXU, ENCFF952JIK | ENCFF299JYV, ENCFF820PHL | |
| TCF7 | ENCFF152RNE | ENCFF512IAI | ENCFF928MIN | |
| TCF7L2 | | ENCFF556FYF | | |
| TEAD4 | | ENCFF547MLB | | |
| TFAP4 | | | ENCFF912SQI | |
| THAP1 | | ENCFF130TPD | | |
| THRA | | ENCFF309DMZ | | |
| THRAP3 | | ENCFF354UUL | | |
| TRIM22 | ENCFF830TFU | | ENCFF063GDN | |
| TRIM24 | | ENCFF063NXI, ENCFF950TOJ | | |
| TRIM28 | | ENCFF168KHS, ENCFF623ELO, ENCFF996AMX | | |
| TRIP13 | | ENCFF534VQL | | |
| U2AF1 | | ENCFF482DRO | ENCFF034KUO | |
| U2AF2 | | ENCFF134HBP | ENCFF562ADR | |
| UBTF | ENCFF295ZLM | ENCFF345RRM, ENCFF403TAF | | |
| USF1 | | | ENCFF914IFQ | |
| USF2 | ENCFF514SWA | ENCFF425FVY | | |
| WRNIP1 | ENCFF514DDI | | | |
| XRCC3 | | ENCFF115PGE | | |
| XRCC5 | | ENCFF929TWP | | |
| YBX1 | ENCFF500RBO | ENCFF520DIY | ENCFF332FUE | |
| YBX3 | | ENCFF508WCC | | |
| YY1 | ENCFF223MUF | ENCFF024TJO, ENCFF635XCI | ENCFF177YDT | ENCFF459TWF |
| ZBED1 | ENCFF630FLK | ENCFF388TYU | | |
| ZBTB11 | | ENCFF913HCQ | | |

Table 1: (continued)

| TF | GM12878 | K562 | HepG2 | liver |
|---|---|---|---|---|
| ZBTB2 | | ENCFF189WAO | | |
| ZBTB33 | ENCFF475DID | ENCFF556STK | ENCFF943WRA | ENCFF727ZIT |
| ZBTB40 | ENCFF084IUW | ENCFF088LZZ | | |
| ZBTB5 | | ENCFF014KUI, ENCFF813GMP | | |
| ZBTB7A | | ENCFF245LRG | ENCFF953JQD | |
| ZBTB8A | | ENCFF328SSL | | |
| ZC3H11A | | ENCFF478PGJ | | |
| ZEB2 | | ENCFF553KIK, ENCFF808NWU | | |
| ZFP36 | ENCFF224WII | | | |
| ZFP91 | | ENCFF150ZBH | | |
| ZHX1 | | ENCFF495BPY | | |
| ZHX2 | | | ENCFF964KDQ | |
| ZKSCAN1 | | ENCFF704VDI | ENCFF721NEC | |
| ZMIZ1 | | ENCFF526PMI | | |
| ZMYM3 | | ENCFF195IFB | ENCFF769SEZ | |
| ZNF143 | ENCFF153TQR | ENCFF700GZI | | |
| ZNF184 | | ENCFF855CUN | | |
| ZNF184A | | ENCFF760EPB | | |
| ZNF207 | ENCFF676BIG | | ENCFF657ZXY | |
| ZNF217 | ENCFF200SLC | | | |
| ZNF24 | ENCFF313HBL | ENCFF007EEV, ENCFF260CBQ, ENCFF723JDW | ENCFF858WPR, ENCFF904QAD | |
| ZNF274 | | ENCFF323AWS, ENCFF498VQZ | | |
| ZNF280A | | ENCFF074WRG | | |
| ZNF282 | | ENCFF596JDS | ENCFF482XNG | |
| ZNF316 | | ENCFF056SEM, ENCFF806GUF | | |
| ZNF318 | | ENCFF082RIZ, ENCFF577LQR | | |
| ZNF384 | ENCFF942MDT | ENCFF106YXG | ENCFF950VAR | |
| ZNF407 | | ENCFF538GSS, ENCFF644XES | | |
| ZNF592 | ENCFF615DTQ | ENCFF972UGK | | |
| ZNF622 | ENCFF777DVJ | | | |
| ZNF639 | | ENCFF008JJE, ENCFF404EVY | | |
| ZNF687 | ENCFF137BRA | | | |

Table 1: (continued)

| TF | GM12878 | K562 | HepG2 | liver |
|---|---|---|---|---|
| ZNF830 | | ENCFF951OSW, ENCFF979NKM | | |
| ZPF36 | | | ENCFF166GKK | |
| ZSCAN29 | ENCFF214NJL | ENCFF908ZLN, ENCFF979GFF | | |
| ZTBTB40 | | | ENCFF624WDI | |
| ZZZ3 | ENCFF260NAX | ENCFF945HJR | | |

Table 2: Summary of the TFs under study. The first column specifies the TF and the following four columns indicate the availability of ChIP-seq data for the different cell types. In column "Methylation", "NA" indicates that data sets have been available only for a single cell type, preventing across cell type predictions; "n" indicates that no significant increase in performance could be observed when including methylation information; "i" indicates inconsistent results between the different strategies for sampling negative training examples; "y" indicates consistent and significant increase in performance when including methylation information. The entries in column "Methyl. & Deps. have the same meaning but for increasing performance when considering methylation information and intra-motif dependencies. In column "Literature", "-", "+" and "s" indicate negative or positive influence of methylation or general methylation sensitivity, respectively; "new" indicates cases that have been identified as methylation sensitive in this study.

| TF | GM12878 | HepG2 | K562 | liver | Methylation | Methyl. & Deps. | Literature |
|---|---|---|---|---|---|---|---|
| AFF1 | | | x | | NA | NA | |
| AGO1 | | x | x | | n | n | |
| AGO2 | | x | | | NA | NA | |
| ARHGAP35 | | | x | | NA | NA | |
| ARID1B | | | x | | NA | NA | |
| ARID2 | | | x | | NA | NA | |
| ARID3A | x | x | x | | y | n | new |
| ARNT | x | x | x | | y | n | - (2) |
| ASH1L | | | x | | NA | NA | |
| ASH2L | x | x | | | n | n | |
| ATF2 | x | x | x | | i | n | - (3) |
| ATF3 | | x | x | x | y | y | - (3) |
| ATF4 | | | x | | NA | NA | - (4; 5) |
| ATF7 | x | x | x | | y | n | - (3) |
| ATM | | x | | | NA | NA | |
| BACH1 | x | | | | NA | NA | |
| BATF | x | | | | NA | NA | different motif (6) |
| BCL11A | x | | | | NA | NA | |
| BCL3 | x | | | | NA | NA | |

Table 2: (continued)

| TF | GM12878 | HepG2 | K562 | liver | Methylation | Methyl. & Deps. | Literature |
|---|---|---|---|---|---|---|---|
| BCLAF1 | x | x | x | | n | n | |
| BCOR | | | x | | NA | NA | |
| BHLHE40 | x | x | x | | y | n | - (3) |
| BMI1 | x | | x | | i | n | |
| BRCA1 | x | x | x | | y | n | new |
| BRD4 | | x | x | | i | n | |
| BRD9 | | | x | | NA | NA | |
| CBFA2T2 | | | x | | NA | NA | |
| CBFA2T3 | | | x | | NA | NA | |
| CBFB | x | | | | NA | NA | |
| CBX1 | | | x | | NA | NA | |
| CBX2 | | x | | | NA | NA | |
| CBX3 | x | | x | | n | n | |
| CBX5 | x | | x | | n | n | |
| CC2D1A | | | x | | NA | NA | |
| CCAR2 | | x | x | | i | n | |
| CDC5L | | | x | | NA | NA | |
| CEBPB | x | x | x | | n | n | +/- (4; 7; 8; 5) |
| CEBPBZ | | | x | | NA | NA | |
| CEBPZ | x | x | | | i | n | |
| CHAMP1 | | | x | | NA | NA | |
| CHD1 | x | | | | NA | NA | |
| CHD2 | x | x | | | i | n | |
| CHD4 | x | x | | | y | n | new |
| COPS2 | | | x | | NA | NA | |
| CREB1 | | x | | | NA | NA | - (3) |
| CREB3L1 | | | x | | NA | NA | |
| CREM | x | x | x | | y | n | - (3) |
| CTBP1 | | | x | | NA | NA | |
| CTCF | x | x | x | | n | n | - (2; 9; 10; 11) |
| CUX1 | x | | x | | i | n | - (3) |
| DACH1 | | | x | | NA | NA | |
| DDX20 | | | x | | NA | NA | |
| DEAF1 | | | x | | NA | NA | |
| DNMT1 | | | x | | NA | NA | |
| DPF2 | x | | x | | n | n | |
| E2F1 | | | x | | NA | NA | - (3) |
| E2F4 | x | | | | NA | NA | - (3) |
| E2F6 | | | x | | NA | NA | |
| E2F7 | | | x | | NA | NA | - (3) |

Table 2: (continued)

| TF | GM12878 | HepG2 | K562 | liver | Methylation | Methyl. & Deps. | Literature |
|---|---|---|---|---|---|---|---|
| E2F8 | x | | x | | i | n | |
| E4F1 | x | | x | | i | n | |
| EBF1 | x | | | | NA | NA | |
| EED | x | | | | NA | NA | |
| EGR1 | | | x | x | n | n | |
| EHMT2 | | x | x | | i | n | |
| EKL1 | x | | | | NA | NA | |
| ELF1 | x | x | x | | y | n | - (4; 3) |
| ELF4 | | | x | | NA | NA | - (3) |
| ELK1 | | | x | | NA | NA | - (4; 3) |
| EP300 | x | x | x | | i | n | |
| EP400 | | | x | | NA | NA | |
| ESRRA | x | | x | | i | n | |
| ETS1 | x | x | x | | i | y | - (4) |
| ETV4 | | x | | | NA | NA | - (3) |
| ETV6 | x | | x | | i | n | |
| EWSR1 | | | x | | NA | NA | |
| EZH2 | x | x | | | n | n | |
| FIP1L1 | | x | x | | n | n | |
| FOSL1 | | | x | | NA | NA | |
| FOSL2 | | x | | | NA | NA | |
| FOXA1 | | x | x | x | y | n | new |
| FOXA2 | | x | | x | y | y | new |
| FOXK2 | x | x | x | | y | n | new |
| FOXM1 | | | x | | NA | NA | |
| FOXP1 | | x | | | NA | NA | |
| FUS | | x | x | | n | n | |
| GABPA | x | x | x | x | y | n | - (3) |
| GABPB1 | | | x | | NA | NA | |
| GATA1 | | | x | | NA | NA | + (4) |
| GATA2 | | | x | | NA | NA | +/- (4; 8) |
| GATA4 | | x | | | NA | NA | + (4) |
| GATAD2A | | | x | | NA | NA | |
| GATAD2B | x | | x | | i | n | |
| GMEB1 | | | x | | NA | NA | - (3) |
| GTF2F1 | | x | x | | i | n | |
| HCFC1 | x | x | x | | y | y | new |
| HDAC1 | | x | x | | n | n | |
| HDAC2 | x | x | x | | y | n | new |
| HDAC3 | | | x | | NA | NA | |
| HDAC6 | | x | x | | n | n | |
| HDGF | x | | x | | n | n | |

Table 2: (continued)

| TF | GM12878 | HepG2 | K562 | liver | Methylation | Methyl. & Deps. | Literature |
|---|---|---|---|---|---|---|---|
| HES1 | | | x | | NA | NA | - (3) |
| HMBOX1 | | | x | | NA | NA | |
| HNF1A | | x | | | NA | NA | |
| HNF4A | | x | | x | y | y | new |
| HNF4G | | x | | x | y | y | new |
| HNRNPH1 | | x | x | | i | n | |
| HNRNPK | | x | x | | n | n | |
| HNRNPL | | x | x | | n | n | |
| HNRNPLL | | x | x | | i | n | |
| HNRNPUL1 | | x | x | | i | n | |
| HSF1 | x | | | | NA | NA | |
| IKZF1 | x | x | x | | i | n | |
| IKZF2 | x | | | | NA | NA | |
| ILF3 | | | x | | NA | NA | |
| IRF2 | | | x | | NA | NA | + (3) |
| IRF3 | x | | | | NA | NA | |
| IRF4 | x | | | | NA | NA | |
| IRF5 | x | | | | NA | NA | |
| JUN | | | x | | NA | NA | - (3) |
| JUNB | x | | x | | i | n | - (3) |
| JUND | x | x | x | x | y | y | - (3) |
| KAT2A | x | | | | NA | NA | |
| KAT2B | | x | | | NA | NA | |
| KAT8 | | | x | | NA | NA | |
| KDM1A | x | x | x | | i | n | |
| KDM4B | | | x | | NA | NA | |
| KDM5A | | x | | | NA | NA | |
| KDM5B | | | x | | NA | NA | |
| KLF16 | | | x | | NA | NA | + (3) |
| KLF5 | x | | | | NA | NA | |
| L3MBTL2 | | | x | | NA | NA | |
| LCORL | | x | | | NA | NA | |
| LEF1 | | | x | | NA | NA | |
| MAFF | | x | x | | n | n | |
| MAFK | x | x | x | | n | n | |
| MAX | x | x | x | x | y | y | s/- (2; 3) |
| MAZ | x | x | | | i | n | |
| MBD2 | | | x | | NA | NA | |
| MCM2 | | | x | | NA | NA | |
| MCM3 | | | x | | NA | NA | |
| MCM5 | | | x | | NA | NA | |
| MCM7 | | | x | | NA | NA | |

Table 2: (continued)

| TF | GM12878 | HepG2 | K562 | liver | Methylation | Methyl. & Deps. | Literature |
|---|---|---|---|---|---|---|---|
| MEF2A | x | | x | | n | n | |
| MEF2B | x | | | | NA | NA | |
| MEF2C | x | | | | NA | NA | |
| MEIS2 | | | x | | NA | NA | |
| MGA | | | x | | NA | NA | |
| MIER1 | | | x | | NA | NA | |
| MITF | | | x | | NA | NA | |
| MLLT1 | x | | x | | n | n | |
| MNT | | x | x | | y | n | s/- (2) |
| MTA1 | | | x | | NA | NA | |
| MTA2 | x | | x | | y | n | new |
| MTA3 | x | | x | | i | n | |
| MXI1 | x | | x | | i | n | |
| MYB | x | | | | NA | NA | |
| MYBL2 | | | x | | NA | NA | - (3) |
| MYC | | | x | | NA | NA | |
| MYNN | | | x | | NA | NA | |
| NBN | x | x | | | n | n | |
| NCOA1 | | | x | | NA | NA | |
| NCOA2 | | | x | | NA | NA | |
| NCOA4 | | | x | | NA | NA | |
| NCOA6 | | | x | | NA | NA | |
| NCOR1 | | x | x | | n | n | |
| NEUROD1 | | | x | | NA | NA | |
| NFATC1 | x | | | | NA | NA | + (3) |
| NFATC3 | x | | x | | y | n | + (3) |
| NFE2 | | | x | | NA | NA | |
| NFE2L2 | | x | | | NA | NA | |
| NFIC | x | | x | | i | n | |
| NFKB | | x | | | NA | NA | |
| NFRKB | | | x | | NA | NA | |
| NFXL1 | x | | x | | n | n | |
| NFYA | x | | | | NA | NA | s/- (2) |
| NFYB | x | | | | NA | NA | |
| NONO | | x | x | | y | y | new |
| NR0B1 | | | x | | NA | NA | |
| NR2C1 | x | | x | | n | n | |
| NR2C2 | x | | x | | y | n | new |
| NR2F1 | x | | x | | n | n | |
| NR2F2 | | | x | x | i | n | |
| NR2F6 | | x | x | | n | n | |
| NR3C1 | | | x | | NA | NA | |

Table 2: (continued)

| TF | GM12878 | HepG2 | K562 | liver | Methylation | Methyl. & Deps. | Literature |
|---|---|---|---|---|---|---|---|
| NRF1 | x | x | x | | y | n | - (2) |
| NUFIP1 | | | x | | NA | NA | |
| PAX5 | x | | | | NA | NA | |
| PAX8 | x | | | | NA | NA | + (3) |
| PBX3 | x | | | | NA | NA | |
| PCBP1 | | x | x | | n | n | |
| PCBP2 | | x | x | | n | n | |
| PHB2 | | x | x | | n | n | |
| PHF20 | | | x | | NA | NA | |
| PHF21A | | | x | | NA | NA | |
| PHF8 | | x | x | | n | n | |
| PKNOX1 | x | | x | | y | n | new |
| PLRG1 | | x | | | NA | NA | |
| PML | | | x | | NA | NA | |
| POU5F1 | | | x | | NA | NA | |
| PRDM10 | | | x | | NA | NA | |
| PRPF4 | | x | x | | n | n | |
| PTBP1 | | x | x | | n | n | |
| PYGO2 | | | x | | NA | NA | |
| RAD21 | x | x | | x | y | y | new |
| RAD51 | x | x | x | | y | y | new |
| RB1 | x | | x | | i | n | |
| RBBP5 | x | | x | | n | n | |
| RBFOX2 | | x | x | | n | n | |
| RBM14 | | | x | | NA | NA | |
| RBM15 | | | x | | NA | NA | |
| RBM17 | | | x | | NA | NA | |
| RBM22 | | x | x | | i | n | |
| RBM25 | | | x | | NA | NA | |
| RBM34 | | | x | | NA | NA | |
| RBM39 | | x | x | | n | n | |
| RCOR1 | x | x | x | | i | n | |
| RELB | x | | | | NA | NA | |
| REST | x | x | x | x | i | n | - (10) |
| RFX1 | | x | x | | n | n | |
| RFX5 | x | x | x | | i | n | |
| RLF | | | x | | NA | NA | |
| RNF2 | | x | x | | y | n | new |
| RUNX1 | | | x | | NA | NA | |
| RUNX3 | x | | | | NA | NA | - (8; 3) |
| RXRA | x | x | | x | i | n | |
| SAFB | | | x | | NA | NA | |

Table 2: (continued)

| TF | GM12878 | HepG2 | K562 | liver | Methylation | Methyl. & Deps. | Literature |
|---|---|---|---|---|---|---|---|
| SAFB2 | | | x | | NA | NA | |
| SAP30 | | | x | | NA | NA | |
| SETDB1 | | | x | | NA | NA | |
| SIN3A | x | x | x | | i | n | |
| SIN3B | | x | x | | i | n | |
| SIX5 | x | | x | | y | n | new |
| SKI | | x | | | NA | NA | |
| SKIL | x | | x | | i | n | |
| SMAD1 | x | | x | | i | n | |
| SMAD2 | | | x | | NA | NA | |
| SMAD5 | x | | x | | n | n | + (3) |
| SMARCA4 | | | x | | NA | NA | |
| SMARCA5 | x | | x | | i | n | |
| SMARCB1 | | | x | | NA | NA | |
| SMARCC2 | | x | x | | n | n | |
| SMARCE1 | | x | x | | n | n | |
| SMC3 | x | x | x | | i | n | |
| SNIP1 | | | x | | NA | NA | |
| SNRNP70 | | x | x | | n | n | |
| SOX13 | | x | | | NA | NA | |
| SOX6 | | x | x | | n | n | |
| SP1 | | x | x | x | y | y | +/- (3; 12; 13; 14) |
| SPI1 | x | | x | | n | n | |
| SREBF1 | | | x | | NA | NA | - (3) |
| SRF | x | | | | NA | NA | |
| SRSF3 | | | x | | NA | NA | |
| SRSF4 | | x | | | NA | NA | |
| SRSF7 | | | x | | NA | NA | |
| SRSF9 | | x | x | | n | n | |
| STAT1 | x | | | | NA | NA | + (4) |
| STAT3 | x | | | | NA | NA | |
| STAT5A | | | x | | NA | NA | + (4) |
| SUPT20H | x | | | | NA | NA | |
| SUPT5H | | | x | | NA | NA | |
| SUZ12 | | x | x | | i | n | |
| SYNCRIP | | x | | | NA | NA | |
| TAF1 | x | x | | x | y | n | new |
| TAF15 | | x | x | | n | n | |
| TAF7 | | | x | | NA | NA | |
| TAF9B | | | x | | NA | NA | |
| TAL1 | | | x | | NA | NA | |

Table 2: (continued)

| TF | GM12878 | HepG2 | K562 | liver | Methylation | Methyl. & Deps. | Literature |
|---|---|---|---|---|---|---|---|
| TARDBP | x | x | x | | i | n | |
| TBL1XR1 | x | x | x | | y | n | new |
| TBP | x | x | x | | n | n | |
| TBX21 | x | | | | NA | NA | |
| TBX3 | | x | | | NA | NA | |
| TCF12 | x | x | x | | n | n | |
| TCF7 | x | x | x | | i | n | |
| TCF7L2 | | | x | | NA | NA | |
| TEAD4 | | | x | | NA | NA | |
| TFAP4 | | x | | | NA | NA | |
| THAP1 | | | x | | NA | NA | |
| THRA | | | x | | NA | NA | |
| THRAP3 | | | x | | NA | NA | |
| TRIM22 | x | x | | | n | n | |
| TRIM24 | | | x | | NA | NA | |
| TRIM28 | | | x | | NA | NA | |
| TRIP13 | | | x | | NA | NA | |
| U2AF1 | | x | x | | n | n | |
| U2AF2 | | x | x | | n | n | |
| UBTF | x | | x | | n | n | |
| USF1 | | x | | | NA | NA | - (2; 3) |
| USF2 | x | | x | | y | y | - (3) |
| WRNIP1 | x | | | | NA | NA | |
| XRCC3 | | | x | | NA | NA | |
| XRCC5 | | | x | | NA | NA | |
| YBX1 | x | x | x | | n | n | |
| YBX3 | | | x | | NA | NA | |
| YY1 | x | x | x | x | y | n | different motif (15) |
| ZBED1 | x | | x | | n | n | - (3) |
| ZBTB11 | | | x | | NA | NA | |
| ZBTB2 | | | x | | NA | NA | - (3) |
| ZBTB33 | x | x | x | x | n | n | + (3; 16; 17; 18) |
| ZBTB40 | x | | x | | y | n | new |
| ZBTB5 | | | x | | NA | NA | |
| ZBTB7A | | x | x | | n | n | - (3) |
| ZBTB8A | | | x | | NA | NA | |
| ZC3H11A | | | x | | NA | NA | |
| ZEB2 | | | x | | NA | NA | |
| ZFP36 | x | | x | | n | n | |
| ZFP91 | | | x | | NA | NA | |

Table 2: (continued)

| TF | GM12878 | HepG2 | K562 | liver | Methylation | Methyl. & Deps. | Literature |
|---|---|---|---|---|---|---|---|
| ZHX1 | | | x | | NA | NA | |
| ZHX2 | | x | | | NA | NA | |
| ZKSCAN1 | | x | x | | i | n | |
| ZMIZ1 | | | x | | NA | NA | |
| ZMYM3 | | x | x | | n | n | |
| ZNF143 | x | | x | | n | n | |
| ZNF184 | | | x | | NA | NA | |
| ZNF184A | | | x | | NA | NA | |
| ZNF207 | x | x | | | n | n | |
| ZNF217 | x | | | | NA | NA | |
| ZNF24 | x | x | x | | i | n | |
| ZNF274 | | | x | | NA | NA | + (3) |
| ZNF280A | | | x | | NA | NA | |
| ZNF282 | | x | x | | n | n | |
| ZNF316 | | | x | | NA | NA | |
| ZNF318 | | | x | | NA | NA | |
| ZNF384 | x | x | x | | i | n | |
| ZNF407 | | | x | | NA | NA | |
| ZNF592 | x | | x | | n | n | |
| ZNF622 | x | | | | NA | NA | |
| ZNF639 | | | x | | NA | NA | |
| ZNF687 | x | | | | NA | NA | |
| ZNF830 | | | x | | NA | NA | |
| ZPF36 | | x | | | NA | NA | |
| ZSCAN29 | x | | x | | i | n | |
| ZTBTB40 | | x | | | NA | NA | |
| ZZZ3 | x | | x | | i | n | |

Table 3: Training data sets for the TFs and models considered in Figure 8 and Supplementary Figures 4 to 13.

| cell type | TF | ENCODE-ID |
|---|---|---|
| GM12878 | NRF1 | ENCFF652BRY |
| HepG2 | ATF3 | ENCFF137OEY |
| HepG2 | BHLHE40 | ENCFF361YXC |
| HepG2 | FOXA2 | ENCFF184NAC |
| HepG2 | HNF4A | ENCFF072CXB |
| HepG2 | MAX | ENCFF140PUO |
| HepG2 | YY1 | ENCFF177YDT |
| K562 | CREM | ENCFF021XJN |
| K562 | ELF1 | ENCFF617ZLL |
| K562 | JUND | ENCFF213EYD |

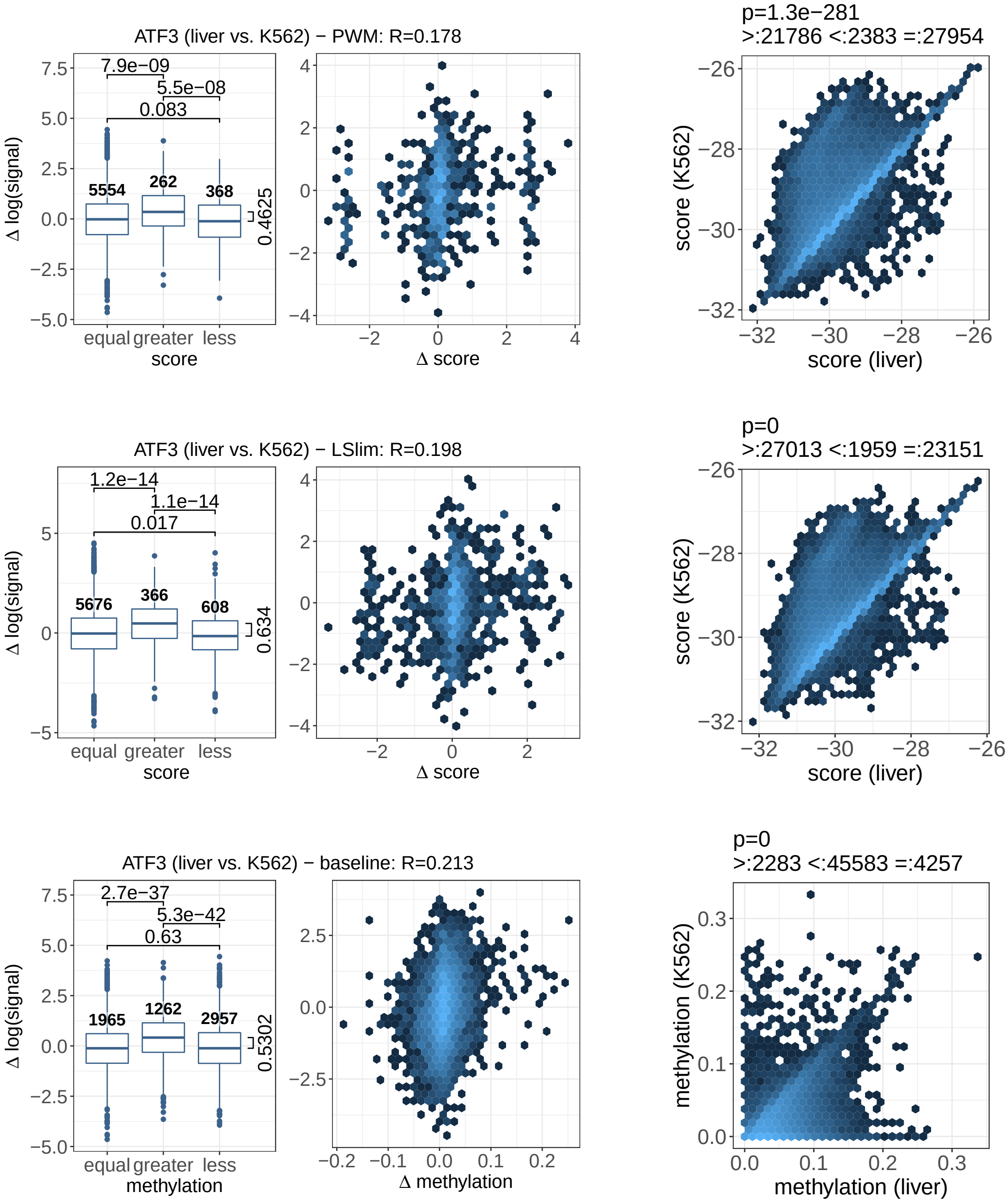


Figure 4: Association between differential model scores and differential binding according to ChIP-seq data for ATF3 in liver and K562 cell types using PWM models, LSlim models, or a baseline model using average methylation levels in a fixed-size region at the peak.

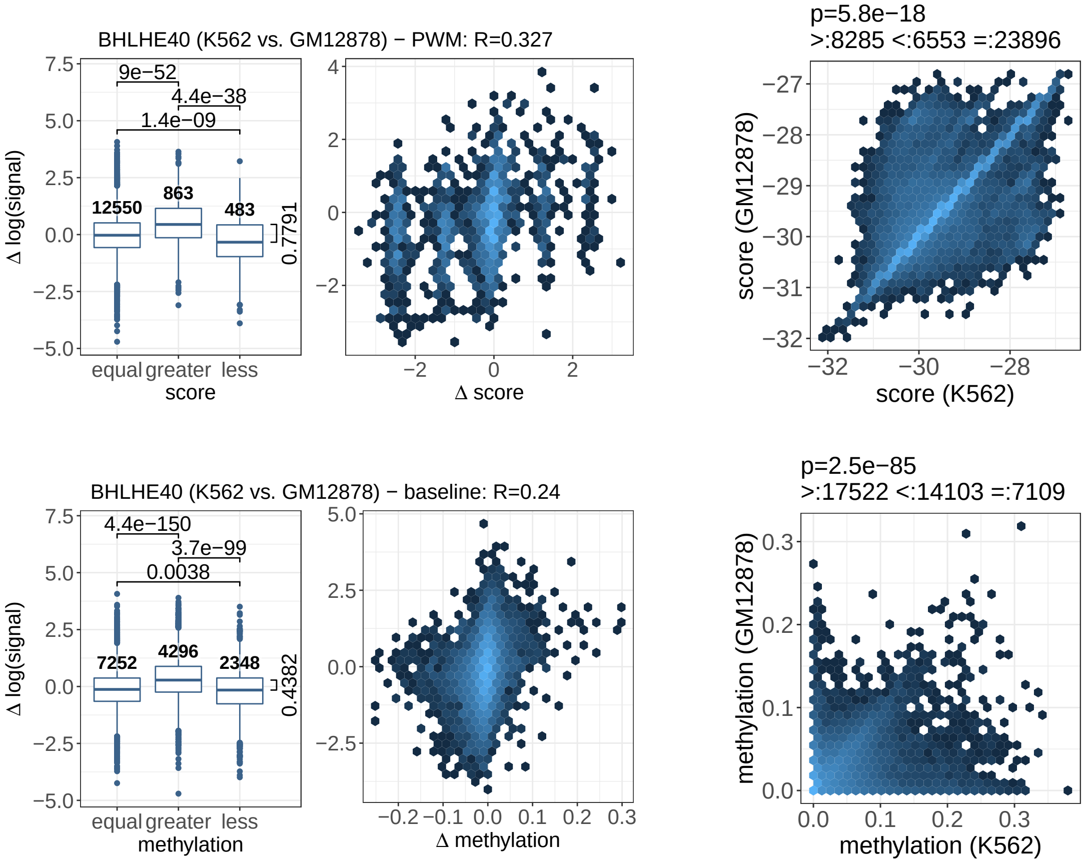


Figure 5: Association between differential model scores and differential binding according to ChIP-seq data for BHLHE40 in K562 and GM12878 cell types using PWM models or a baseline model using average methylation levels in a fixed-size region at the peak.

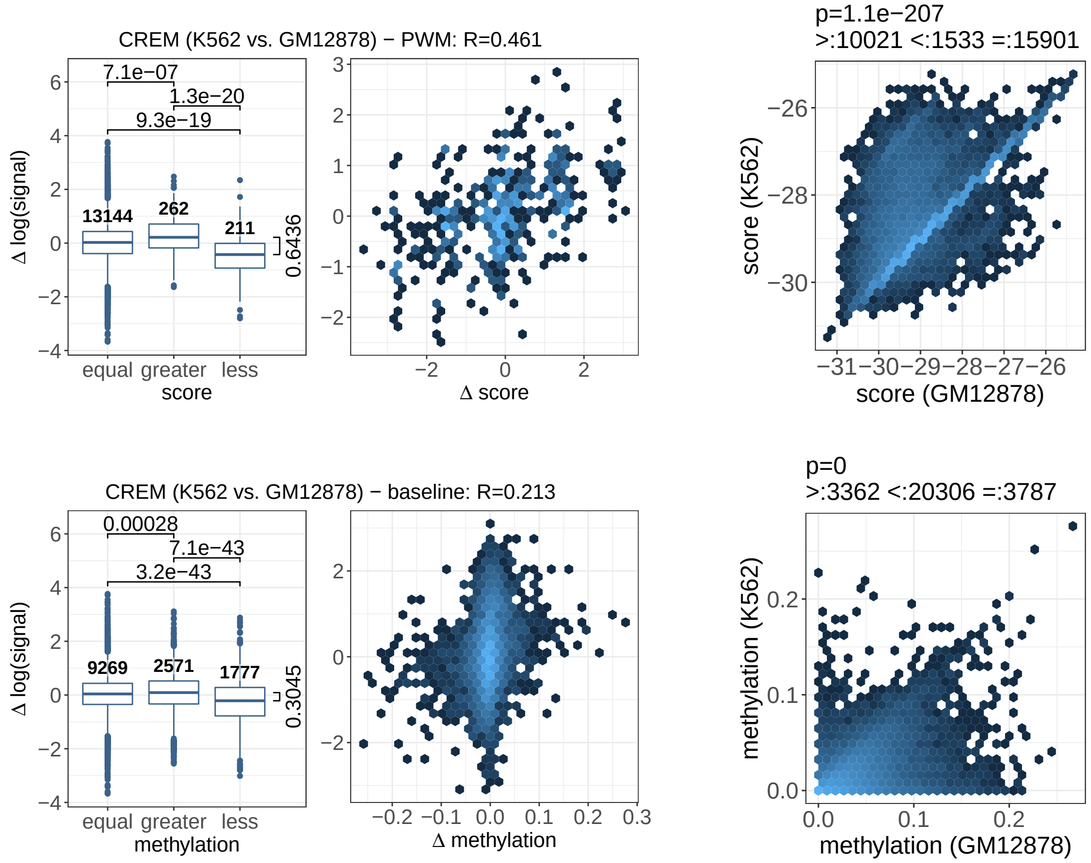


Figure 6: Association between differential model scores and differential binding according to ChIP-seq data for CREM in K562 and GM12878 cell types using PWM models or a baseline model using average methylation levels in a fixed-size region at the peak.

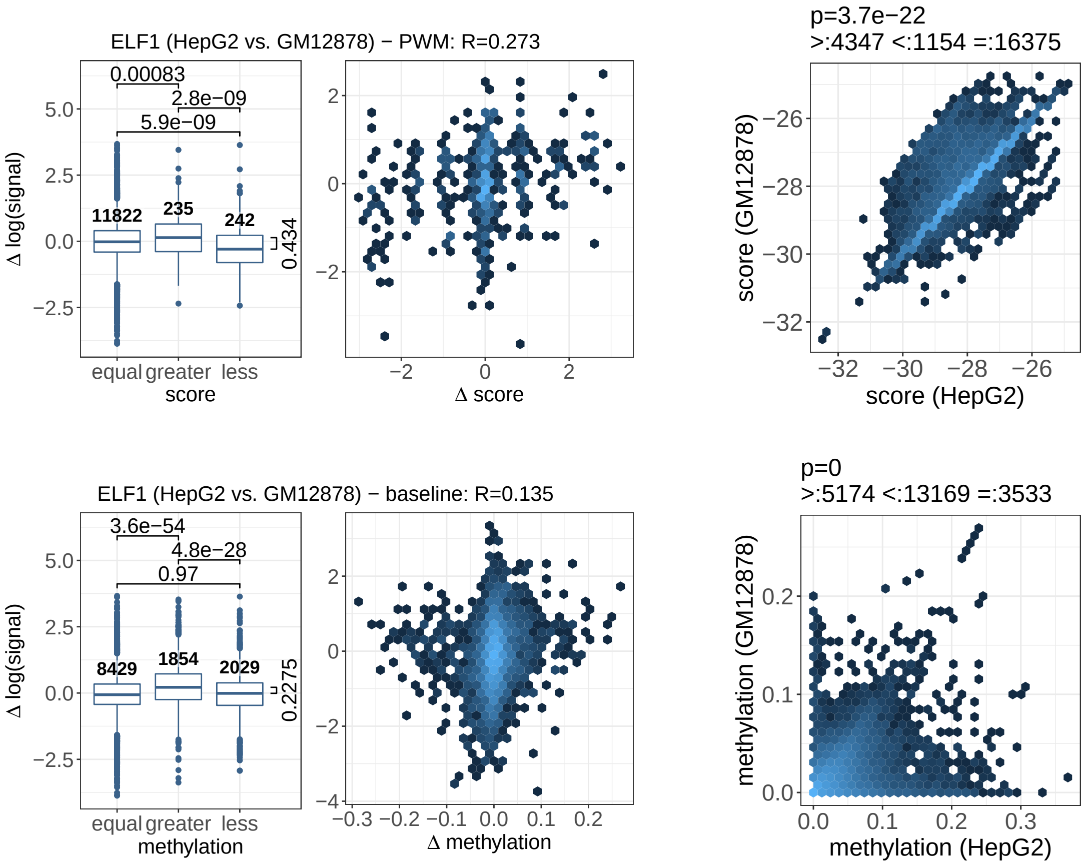


Figure 7: Association between differential model scores and differential binding according to ChIP-seq data for ELF1 in HepG2 and GM12878 cell types using PWM models or a baseline model using average methylation levels in a fixed-size region at the peak.

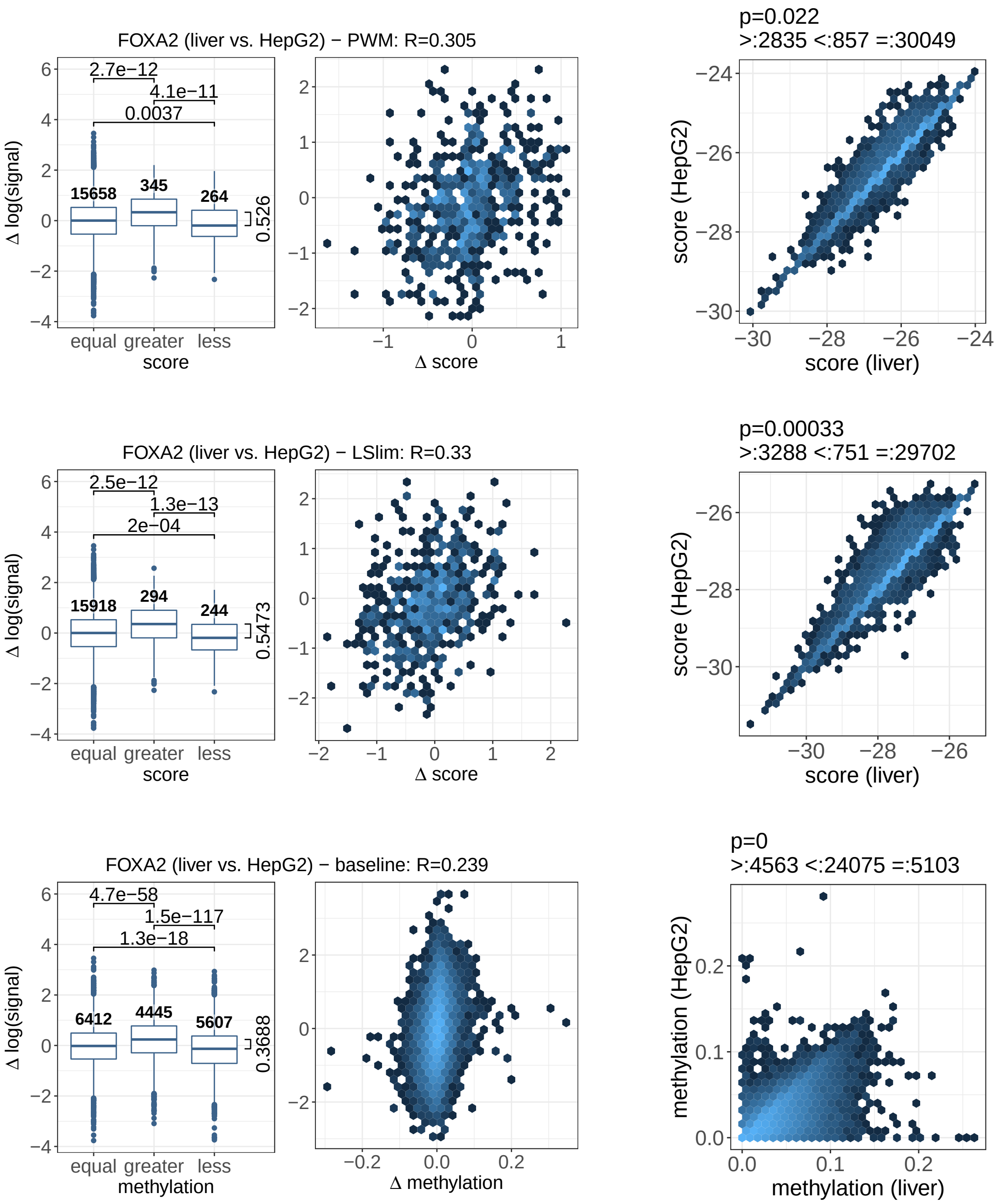


Figure 8: Association between differential model scores and differential binding according to ChIP-seq data for FOXA2 in HepG2 and liver cell types using PWM models, LSlim models, or a baseline model using average methylation levels in a fixed-size region at the peak.

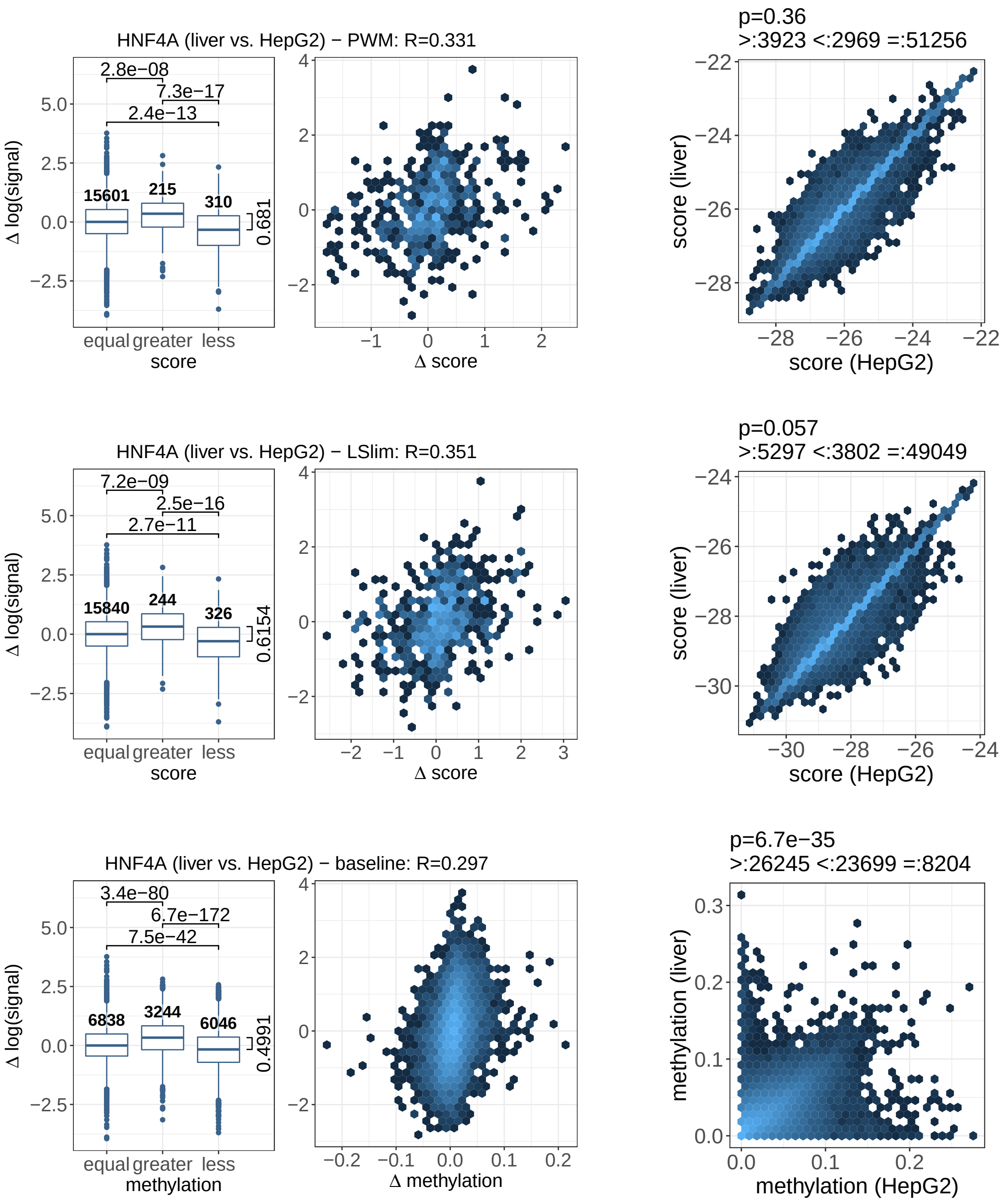


Figure 9: Association between differential model scores and differential binding according to ChIP-seq data for HNF4A in liver and HepG2 cell types using PWM models, LSlim models, or a baseline model using average methylation levels in a fixed-size region at the peak.

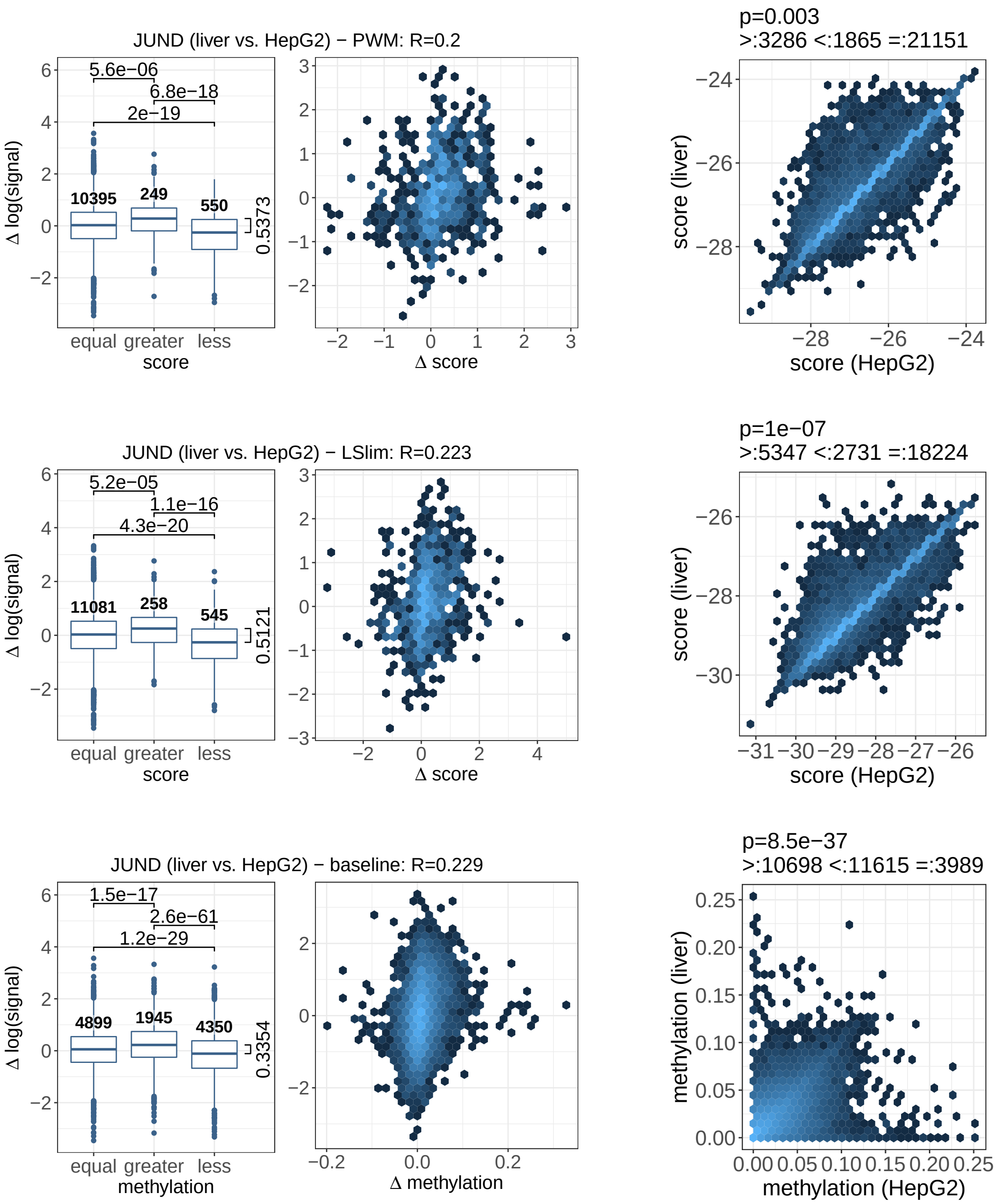


Figure 10: Association between differential model scores and differential binding according to ChIP-seq data for JUND in liver and HepG2 cell types using PWM models, LSlim models, or a baseline model using average methylation levels in a fixed-size region at the peak.

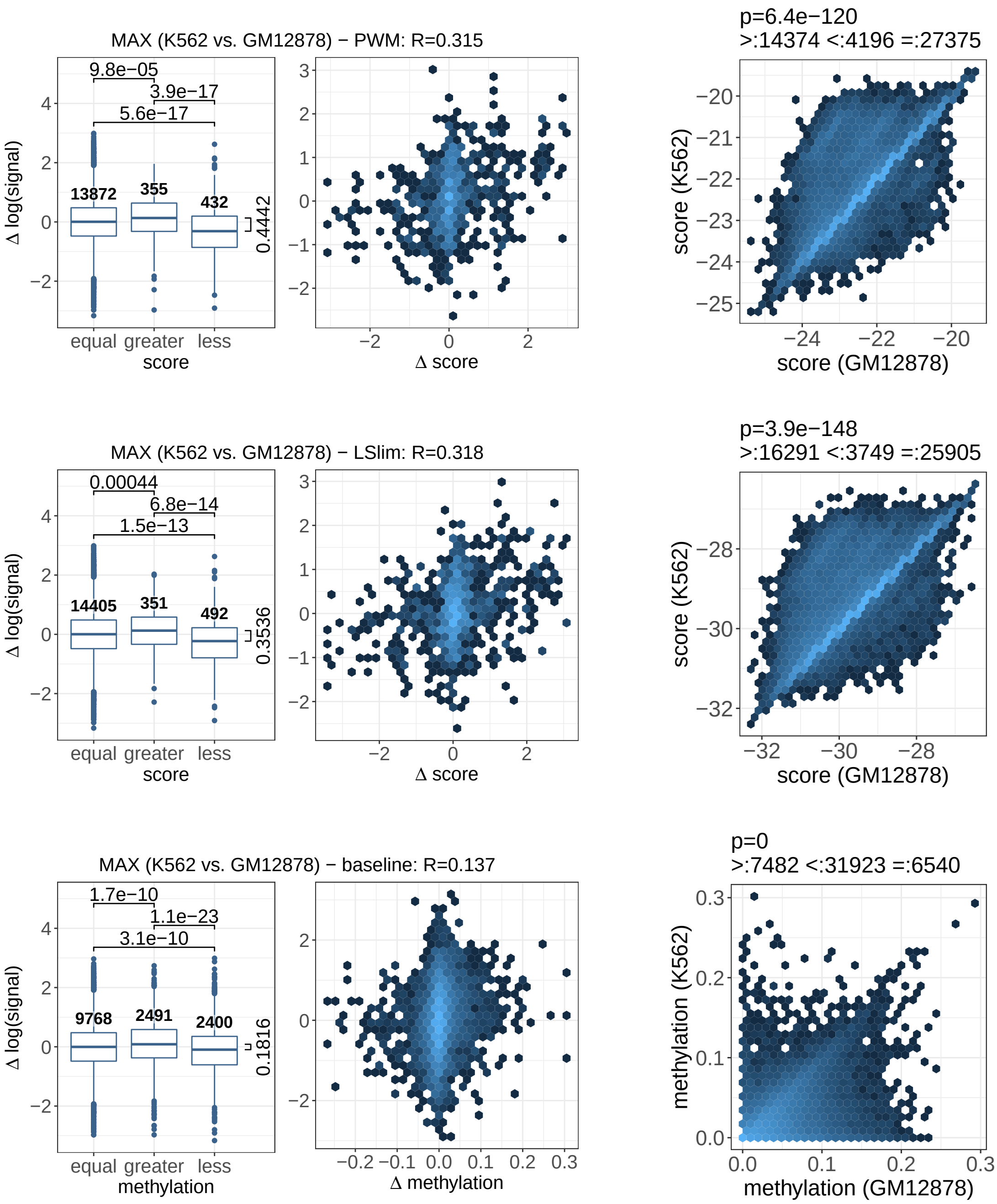


Figure 11: Association between differential model scores and differential binding according to ChIP-seq data for MAX in K562 and GM12878 cell types using PWM models, LSlim models, or a baseline model using average methylation levels in a fixed-size region at the peak.

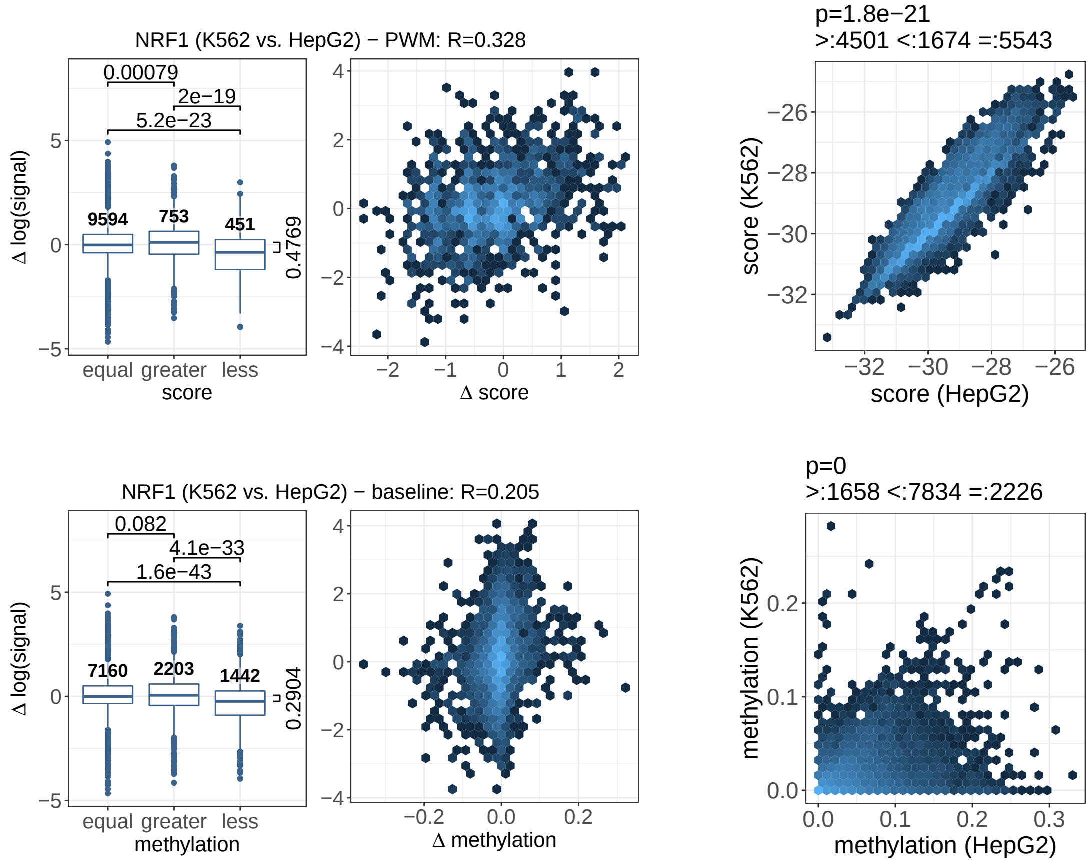


Figure 12: Association between differential model scores and differential binding according to ChIP-seq data for NRF1 in K562 and HepG2 cell types using PWM models or a baseline model using average methylation levels in a fixed-size region at the peak.

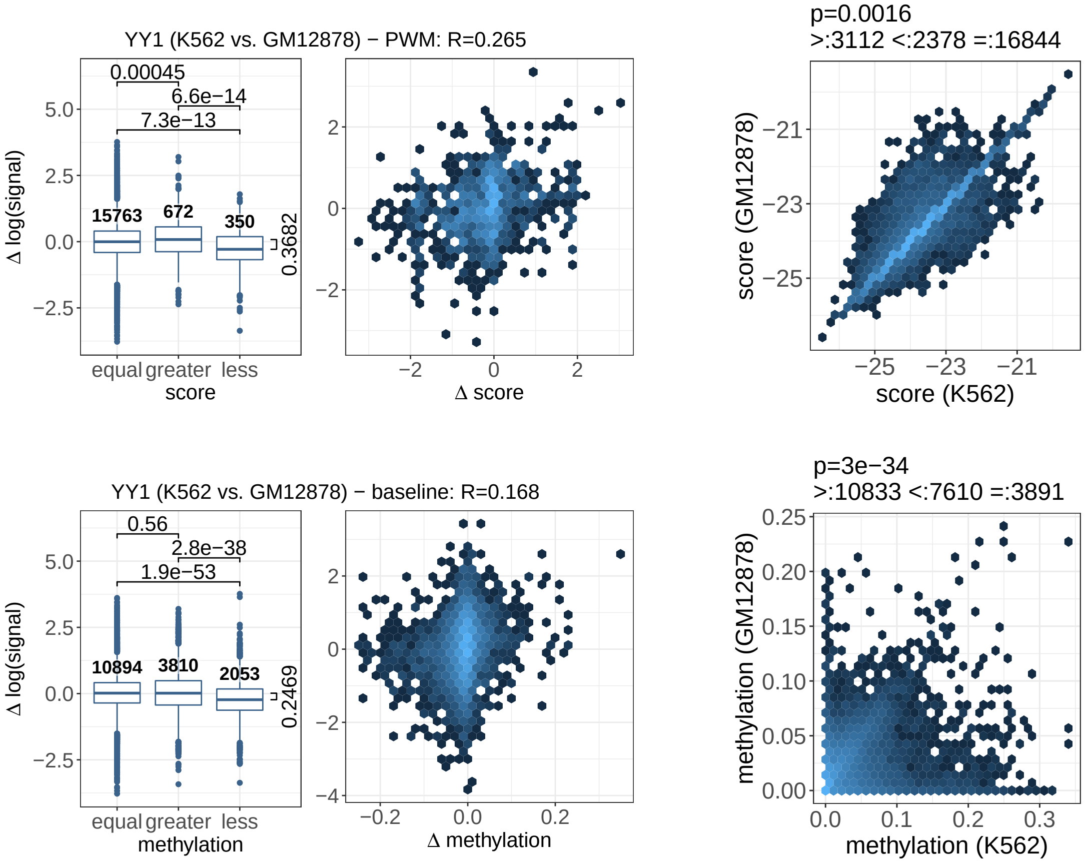


Figure 13: Association between differential model scores and differential binding according to ChIP-seq data for YY1 in K562 and GM12878 cell types using PWM models or a baseline model using average methylation levels in a fixed-size region at the peak.
